# Unravelling the processes controlling pollen formation and functions with cross-species comparative analysis

**DOI:** 10.64898/2026.02.15.705993

**Authors:** Jooa Moon, Eugene Koh, Zhenping Sun, Wenyu Pei, Shilin Du, Zixin Wang, Rohan Shawn Sunil, Jhing Yein Tan, Tay Jie Hao, Tang Chee Chung Kenneth, Boon Chuan Ho, Lily Chen, Nam-Joon Cho, Yi Zhang, Marek Mutwil

## Abstract

Pollen and anthers are essential for plant reproduction, yet the genetic mechanisms governing their formation and function remain incompletely understood. In this study, we performed a comprehensive cross-species comparative analysis leveraging 16,904 RNA-seq samples from 90 plant species, encompassing diverse lineages from bryophytes to angiosperms. By calculating tissue specificity metrics and applying clustering and functional enrichment analyses, we identified both conserved and lineage-specific gene families associated with pollen and anther development. Our computational pipeline allowed for the discovery of previously unstudied genes and predictions of their critical biological roles, which were experimentally validated through mutant characterization, genotyping, and detailed morphological analysis. Notably, we observed evolutionary patterns in pollen morphology, cell wall structure, cytoplasmic content, and aperture characteristics, highlighting differences and gradual transitions among major plant clades. These findings provide significant insights into the evolutionary processes shaping pollen and anther traits, and establish a valuable resource for future studies in plant reproductive biology and biotechnology.

## Introduction

In seed plants, pollen grains – essentially microscopic carriers of the male gametes – are produced in the anthers and form a vital link in the sexual reproductive cycle. Successful development and dispersal of pollen is not only fundamental for plant fertility and species survival, but also directly influences ecosystem biodiversity and crop productivity.^1^ Notably, environmental stresses that disrupt pollen formation can cause male sterility and severe losses in yield (for example, heat-induced pollen abortion in crops like maize can significantly reduce seed set).^2^ The formation of viable pollen within the anther is a complex developmental process involving multiple specialized cell layers (such as the tapetum that nourishes developing microspores) and tightly coordinated gene expression programs.^3^ Unraveling the mechanisms governing pollen and anther development is therefore a longstanding goal in plant biology, with important implications for agriculture (e.g. improving crop fertility and controlling male sterility in hybrid breeding).^4^

Despite their importance, the genetic pathways controlling pollen and anther development remain incompletely understood. Classical genetic studies in model plants like *Arabidopsis thaliana (A. thaliana)* and rice have identified numerous regulatory genes required for anther cell differentiation and pollen maturation^5,6^, and high-throughput transcriptome analyses and genetic screens have catalogued hundreds of pollen-or anther-enriched genes in individual species.^7^ However, these single-species approaches provide only a fragmentary view of the underlying biology. Many genes expressed during microsporogenesis and pollen maturation are still uncharacterized, and at the same time, many uncharacterized genes tend to be specifically expressed in pollen.^8^ Furthermore, it is unclear which are universally important versus specific to certain lineages. Consequently, identifying the full repertoire of pollen/anther-specific genes and deciphering their functions is essential for a complete understanding of plant reproduction and for efforts to manipulate male fertility.^9–11^ Yet achieving such a comprehensive picture with conventional approaches remains challenging, as most functional studies are restricted to a few genetic models, gene family redundancy can mask loss-of-function phenotypes, and large fractions of reproduction-enriched transcripts remain poorly annotated—together limiting both discovery and evolutionary generalization.

Moreover, fostering sustainable environmental and engineering applications based on pollen requires a comparative perspective that spans diverse plant species. At the same time, pollen has emerged as a promising bio-derived engineering material, distinguished by its hierarchical microarchitecture, sporopollenin-based exine, and exceptional resistance to chemical, thermal, and mechanical stress.^12–14^ These properties, originally evolved to ensure reproductive success under harsh environmental conditions, have enabled the transformation of pollen into functional materials such as microcapsules, porous scaffolds, absorbents, and delivery vehicles for applications in biomedicine, environmental remediation, and sustainable manufacturing.^15^ However, most current pollen engineering approaches treat pollen as a passive starting material, with limited consideration of the developmental and genetic programs that give rise to its structure and material properties. Integrating genetic and developmental insights across multiple species is therefore essential to unlock the full potential of pollen as a sustainable, bio-derived material.

One promising strategy to overcome these limitations is cross-species comparative analysis, which leverages the growing wealth of plant genomic and transcriptomic data across diverse lineages.^16–20^ By integrating gene expression profiles from multiple species, comparative transcriptomics can reveal which genes have consistently organ-or tissue-specific roles throughout evolution, and thus highlight core components required for the development and function of the tissue of interest.^8,19,21^ At the same time, this approach can pinpoint lineage-specific genes or pathways, shedding light on how different plant groups (for example, monocots versus eudicots) have evolved distinct mechanisms and traits in pollen development. Indeed, a transcriptomic comparison of ten diverse plant species identified hundreds of organ-and gamete-specific gene families, illustrating the power of broad evolutionary analyses. With large-scale RNA sequencing datasets now available for numerous plant species, extending such cross-species analyses to a wider phylogenetic scope promises to illuminate new genetic factors and evolutionary innovations underlying pollen and anther biology.^16^

In this study, we present a comprehensive comparative transcriptome analysis of spore, pollen and anther development across ∼90 plant species, spanning early-diverging bryophytes and gymnosperms through to angiosperms. We assembled an expression atlas from public datasets and new RNA-seq profiles, and applied a tissue-specificity metric to systematically identify pollen-specific and anther-specific genes in each species. By clustering orthologous genes into families, we could detect both widely conserved gene families associated with male reproduction and lineage-restricted genes unique to particular clades. Our cross-species approach reveals that pollen and anthers possess highly distinctive transcriptomes enriched in specialized functions: notably, we identify numerous conserved pollen gene families. including several previously unstudied genes. as strong candidates for essential roles in pollen formation and function. We also observe clear differences between monocots and dicots in their pollen gene expression profiles, which correlate with known evolutionary divergences in pollen morphology (such as variation in pollen wall structure and aperture traits between these clades). Finally, to validate the biological significance of our findings, we performed targeted loss-of-function experiments in *A.thaliana* for a subset of the conserved pollen/anther-specific genes identified by our pipeline. Disruption of many of these genes led to defects in pollen tube growth, confirming their roles in male reproduction. Together, our work provides new insights into the conserved genetic programs and evolutionary processes that govern pollen formation, and establishes a broad comparative framework and resource for further exploration of plant reproductive biology.

## Results

### Collection of anther-and pollen-specific gene expression data

To enable a systematic comparison of anther-and pollen-enriched transcriptomes across plant lineages, we established a cross-species analysis pipeline that (i) quantifies organ specificity for each gene using the specificity measure (SPM), (ii) aggregates candidates into conserved gene families, and (iii) prioritizes conserved, organ-specific candidates for functional testing using loss-of-function mutants (Fig. 1A). This framework was designed to identify both known and previously uncharacterized genes with high anther or pollen specificity, and to connect these candidates to conserved gene families for downstream comparative and experimental analyses.

**Figure 1.**
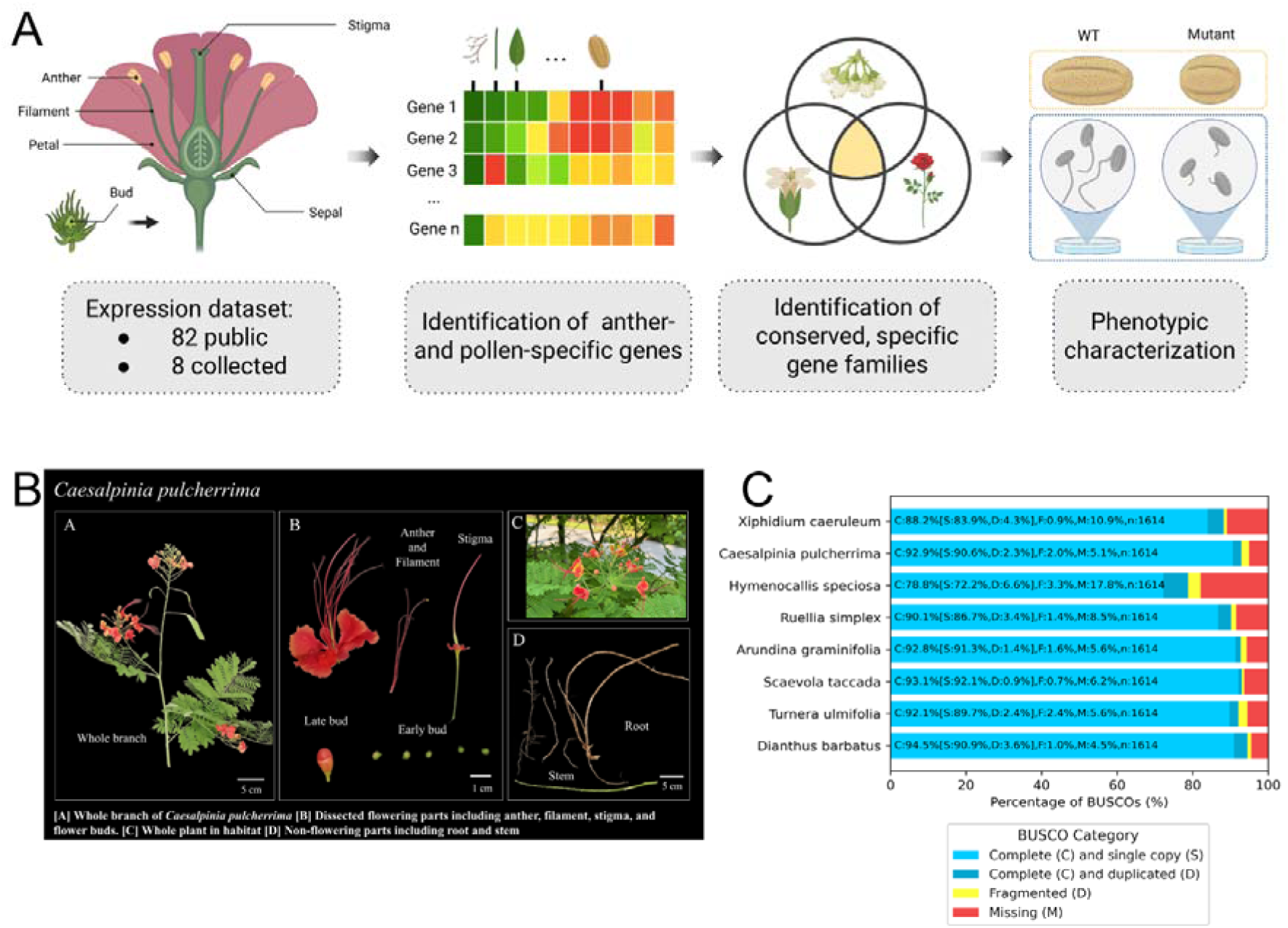
Overview of the in-house sampling and cross-species analysis pipeline. (A) Schematic of the workflow. RNA-seq datasets from public resources and newly generated in-house transcriptomes are integrated to quantify organ-specific expression using the specificity measure (SPM), identify anther-and pollen-specific genes, cluster them into conserved gene families across species, and prioritize candidates for functional validation via reverse genetics and phenotyping. (B) Representative dissection and sampled organs for *Caesalpinia pulcherrima* collected in Singapore. Reproductive tissues included anther, filament, stigma, and floral buds; vegetative tissues included stem and root. Anther were sampled for all in-house species. (C) BUSCO completeness assessment of de novo assembled transcriptomes from eight in-house species representing five dicots (*C. pulcherrima, Ruellia simplex, Turnera ulmifolia, Dianthus barbatus,* and *Scaevola taccada*) and three monocots (*Hymenocallis speciosa, Xiphidium caeruleum,* and *Arundina graminifolia*). Bars indicate the proportion of complete (single-copy/duplicated), fragmented, and missing BUSCOs.

To generate a broad and well-sampled expression compendium, we integrated RNA-seq datasets from 82 plant species obtained from public repositories (Table S1) with newly generated RNA-seq data from eight species sampled in Singapore (Table S2). For each in-house species, we performed on-site dissections to collect reproductive tissues (including anthers, filaments, stigma, and buds) alongside vegetative tissues (stem, leaf, and root), ensuring that all newly sampled species included anther material (Fig. 1B; Fig. S1). The resulting RNA-seq data were then incorporated into the overall expression dataset used for organ-level specificity profiling.

Because reference genomes were unavailable for the in-house species, we reconstructed their coding sequence resources using a de novo transcriptome assembly workflow (Fig. S2). Assembly quality and completeness were assessed with BUSCO, which indicated consistently high completeness across most transcriptomes (>85% complete BUSCOs), with the exception of *H. speciosa*, which showed 78.8% completeness (Fig. 1C; Table S3; Supplemental Data 1; Supplemental Data 2). Together, these in-house datasets (five dicots and three monocots; Fig. 1C) complement the public compendium and provide the foundation for the comparative identification of conserved anther-and pollen-specific genes across 90 species.

### Identification of pollen-and anther-specific genes in 90 land plants

To robustly identify pollen-and anther-specific genes, we first required a broad baseline of gene expression across multiple samples (defined as cell types, tissues and organs), because specificity can only be interpreted relative to expression in non-reproductive samples. We therefore collated RNA-seq datasets for 90 plant species spanning major land plant lineages and consolidated samples into ten major sample types (microspore, pollen, pollen tube cell, anther, leaf, stem, root, flower, seed, and fruit) using Plant Ontology (PO) terms and their hierarchical relationships (Fig. 2A; Fig. S3). To ensure comparability across studies and species, we removed low-quality RNA-seq datasets based on sequencing depth and alignment statistics (Fig. S4). The resulting compendium comprised of 16,904 annotated RNA-seq samples, providing a broad organ coverage across the phylogeny, with the largest representation coming from leaf and root samples (Fig. 2A). Importantly, reproductive tissues were well represented: pollen RNA-seq data were available for 44 species and anther RNA-seq data for 58 species (Fig. 2A), enabling systematic detection of genes preferentially expressed in male reproductive samples while controlling for expression in other samples.

**Figure 2.**
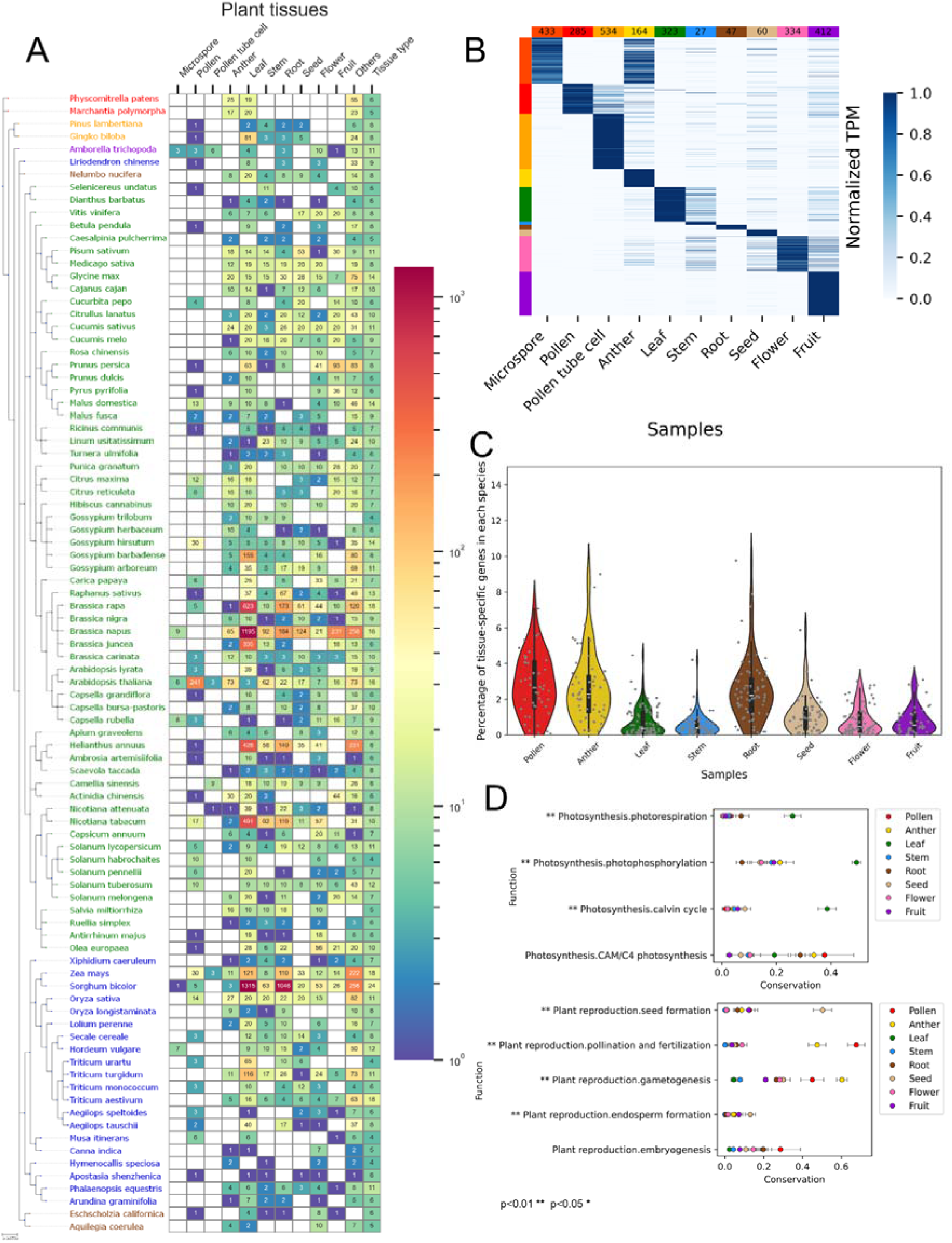
Cross-species RNA-seq compendium and identification of pollen-and anther-specific genes. (A) Scale and composition of the dataset. Heatmap shows the number of RNA-seq samples per major organ category (columns) for each of 90 plant species (rows), ordered by phylogeny. Species names are color-coded by major lineage: bryophytes (red), gymnosperms (orange), ANA grade (violet), basal eudicots (brown), dicots (green), and monocots (blue). Samples were mapped to organ categories using Plant Ontology (PO) terms, consolidated into ten major organs (microspore, pollen, pollen tube cell, anther, leaf, stem, root, flower, seed, fruit). “Others” denotes samples assigned to PO terms outside these categories or too broad/too specific for consolidation. “Tissue Type” refers to the total number of different tissues including both major and non-major organs. (B) Validation of organ specificity calling in *A. thaliana*. Heatmap shows normalized expression profiles across samples for genes classified as sample-specific using the specificity measure (SPM; SPM ≥ 0.8). Rows are genes, columns are samples; genes are grouped by the sample in which they show maximal specificity. (C) Distribution of tissue-specific genes across samples and species. Points represent the proportion of genes per species classified as tissue-specific (SPM ≥ 0.8) for each sample; violin plots show the overall distribution across species and boxplots indicate median and interquartile range. (D) Functional coherence and conservation of sample-specific gene sets. For each species, sample-specific genes were tested for enrichment in curated photosynthesis (upper) and reproduction (lower) functional categories; conservation is shown as the fraction of species in which a category was significantly enriched. Asterisks indicate categories significantly enriched (permutation test (n=10,000)) in the corresponding organ-specific gene sets.

We next quantified sample specificity using the Specificity Measure (SPM)^8^, which captures how strongly a gene’s expression is concentrated in a given sample relative to the full sample compendium for that species. As an initial validation that SPM recapitulates an expected tissue-restricted expression patterns, we applied the approach to *A. thaliana* and visualized organ-specific gene sets across major organs. The resulting profiles showed clear organ-restricted expression blocks consistent with expected tissue specialization (Fig. 2B). Using our predefined and stringent specificity threshold (SPM ≥ 0.8, i.e., 80% of the genes must be sample specific), we identified 285 pollen-specific genes and 164 anther-specific genes in *A. thaliana* (Fig. 2B), supporting that SPM reliably enriches for genes with stringent organ-preferential expression.

We then calculated SPM values for all genes across all species in the dataset (Supplemental Data 3). To compare the extent of tissue specialization across organs, we summarized, for each species, the proportion of genes classified as sample-specific (SPM ≥ 0.8) within each major sample category. Across the 90 species, pollen, anther, root, and seed repeatedly exhibited the highest fractions of sample-specific genes (Fig. 2C), consistent with these organs having particularly specialized transcriptomes.

Finally, to test whether sample-specific gene sets reflect biologically coherent functions across species, we performed pathway enrichment analyses and quantified functional conservation as the fraction of species in which a given pathway is significantly enriched among sample-specific genes. Leaf-specific genes showed strong, consistent enrichment for canonical photosynthesis-related functions—including photorespiration, photophosphorylation, and the Calvin cycle—whereas CAM/C4-related functions were comparatively less leaf-specific across species (Fig. 2D, upper). Within reproductive functions, seed-specific genes were most enriched for seed formation pathways, whereas pollen-and anther-specific genes were most strongly associated with pollination and fertilization-related processes, followed by gametogenesis-associated functions (Fig. 2D, lower). Together, these results indicate that our sample definitions (Fig. S3), quality filtering (Fig. S4), and SPM-based specificity calling (Supplemental Data 3) recover biologically meaningful, sample-relevant programs across diverse plant lineages and provide a foundation for identifying conserved and lineage-specific pollen/anther gene families in subsequent analyses.

### Identification of conserved pollen-and anther-specific genes related to pollen development and function

Motivated by observations that conserved genes often underpin essential biological functions,^22,23^ we hypothesized that pollen-(and anther-) specific genes that recur across many species would be enriched for essential regulators and effectors of male reproduction. To identify these essential genes, we clustered genes into cross-species gene families and quantified, for each family, in how many species at least one member is pollen-specific (SPM ≥ 0.8), using *A. thaliana* as an illustrative reference (Fig. 3A; Table S4,S5).

**Figure 3.**
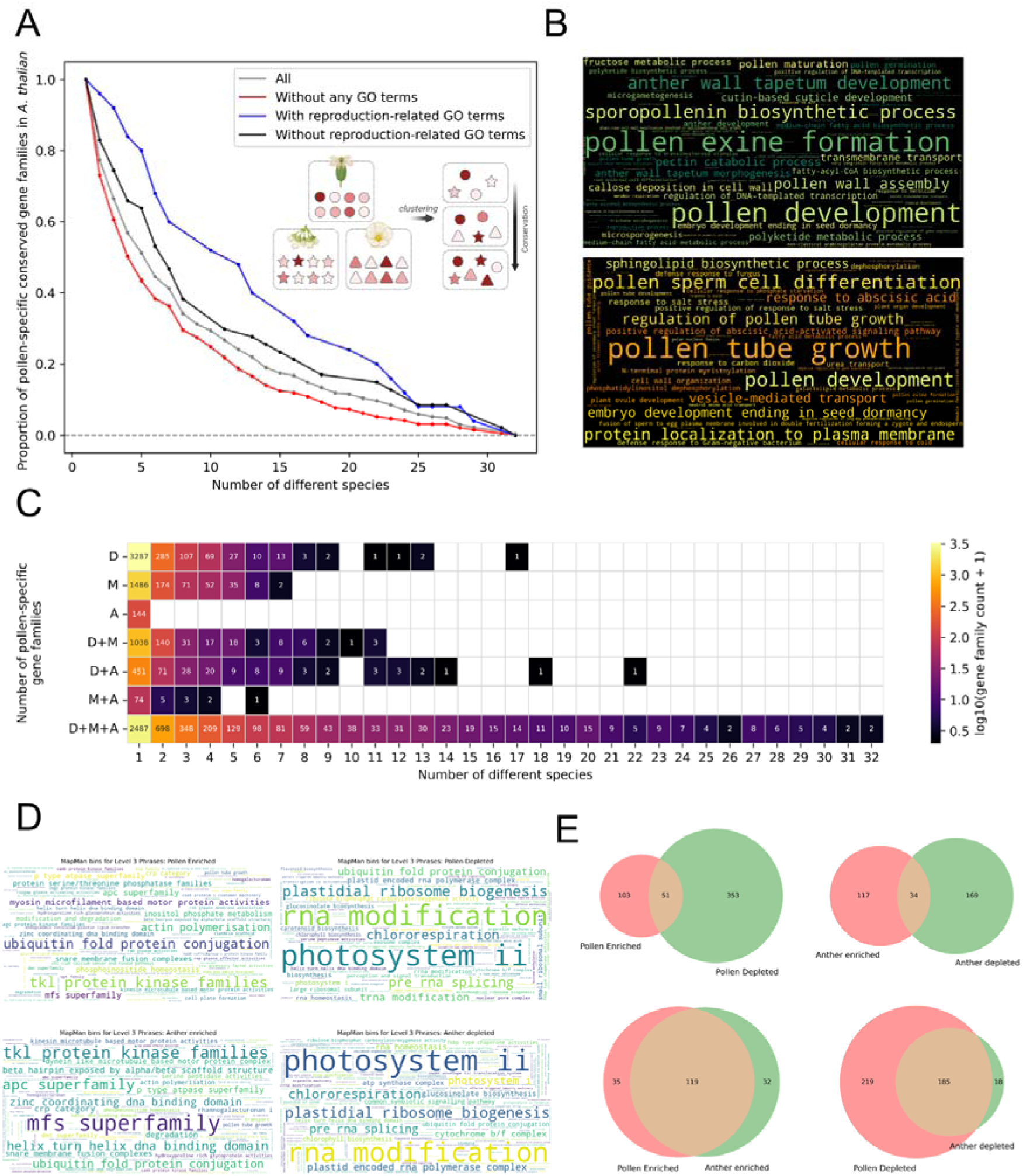
Discovery of the conserved pollen-and anther-specific gene families. (A) Proportion of total and experimentally unstudied conserved pollen-specific gene families (SPM ≥ 0.8) in *A. thaliana*. Number of different species denotes the number of species that has at least one pollen-specific gene within a clustered pollen-specific gene family. A color intensity in the illustrative figure represents SPM, with higher SPM in darker color. (B) Functional annotation of conserved anther-(top) and pollen-(bottom) specific genes with experimental evidence. Gene Ontology(GO) terms were categorized into tissue-associated biological pathways. (C) Number of pollen-specific gene families conserved across major plant clades: Dicots, Monocots and Ancestral lineages (including bryophytes, ANA grade angiosperms, and basal eudicots). (D) Show pollen and anther-enriched and-depleted MapMan functions, while (E) indicates the overlap of these functions in anther and pollen.

As expected, the number of conserved pollen-specific gene families decreased as the conservation requirement became more stringent (Fig. 3A). This monotonic drop is likely due to pollen-associated functions being lineage-restricted or diversified after major evolutionary splits. Despite this expected decline, a substantial set of pollen-specific gene families remained conserved across multiple species, providing a focused candidate set for core pollen biology. When stratifying families by functional annotation, we observed that reproduction-related genes (blue) were, on average, retained as pollen-specific across more species than genes lacking reproduction-associated annotations (Fig. 3A, non-blue lines), consistent with a more conserved “core” male-reproductive program. Notably, among the most conserved pollen-specific gene families, we identified multiple genes that remain experimentally uncharacterized, highlighting a tractable set of high-priority candidates for functional study (Fig. 3A; Table S4). These overall trends were mirrored for anther-specific gene families (Fig. S5A; Table S5), supporting that conservation of organ-specificity is a general feature of male reproductive gene programs.

To connect these conserved, organ-specific gene sets to biological processes, we next examined functional frequency among conserved anther-and pollen-specific genes with experimental evidence. Consistent with the division of labor between tissues, the functions of conserved pollen-specific genes were more specific to processes related to pollen tube growth, guidance, and fertilization, whereas conserved anther-specific genes showed stronger enrichment for pathways associated with pollen formation and structural development (e.g., pollen wall/exine-related processes) (Fig. 3B). This functional partitioning aligns with anthers acting as the developmental environment that produces and supports microspores and maturing pollen, while mature pollen executes the delivery program: rapid germination and directional tube growth culminating in sperm cell delivery to the ovule.

To examine how these conserved gene families are distributed across major plant lineages, we analyzed conservation patterns of pollen-specific families (Fig. 3C) and anther-specific families (Fig. S5B) as a function of their presence in dicots (D), monocots (M), and ancestral lineages (A; bryophytes, gymnosperms, ANA grade, and basal eudicots). For both pollen and anther, the majority of highly conserved gene families were shared across D, M, and A (D+M+A in Fig. 3C and Fig. S5B), indicating that genes essential for male reproduction tend to be maintained across deep evolutionary time rather than being restricted to a single clade. Nevertheless, a subset of families displayed lineage-restricted conservation, being confined to dicots (D), monocots (M), or ancestral lineages (A), or shared only between two of these groups (D+M, D+A, M+A). These lineage-restricted modules likely contribute to clade-specific innovations in pollen and anther biology while the D+M+A classes represent the conserved core.

We then asked which biological functions are preferentially enriched or depleted in pollen and anther relative to non-reproductive genes, using Level 3 MapMan functional categories (Fig. 3D). Across both tissues, photosynthesis-related categories (e.g., “photosystem II,” “plastidial ribosome biogenesis”) and RNA–processing/ RNA-modification functions were markedly depleted in both enriched and depleted sets, consistent with the differentiation of proplastids into amyloplasts and specialized transcriptional programs of male gametophytic tissues.^24,25^ In contrast, categories associated with protein turnover and signaling—such as ubiquitin-related pathways and tyrosine kinase–like (TKL) protein kinases—were significantly enriched in both pollen-and anther-enriched gene sets, highlighting the importance of regulated protein modification and signaling cascades for male gametophyte development and function.^26,27^ Notably, the major facilitator superfamily (MFS) category showed a stronger enrichment in anther than in pollen, suggesting a prominent role for diverse transport processes in supporting pollen development within the anther.^28^

Finally, to quantify functional similarities and differences between pollen and anther, we compared overlaps of enriched and depleted MapMan functions within and between the two tissues (Fig. 3E). Within a given tissue, the overlap between enriched and depleted categories was minimal relative to the total numbers of enriched-only and depleted-only functions, indicating that our classification effectively separates functions preferentially engaged versus downweighted in male reproductive tissues. In contrast, there was substantial overlap between pollen-enriched and anther-enriched categories, and likewise between pollen-depleted and anther-depleted categories (Fig. 3E), indicating that pollen and anther share a largely common functional landscape, potentially caused by pollen being contained within anthers. Together, these analyses show that conserved pollen-and anther-specific gene families are not only deeply shared across land plants, but also occupy a coherent, reproduction-focused functional space dominated by regulatory, signaling, and transport processes.

### Comparative co-expression network analysis reveals conserved transcriptional modules underpinning pollen development

Because co-expression patterns can reveal functionally linked genes within pathways, we next asked whether we could use gene co-expression networks to uncover additional, previously unannotated gene families involved in pollen development. To this end, we constructed anther-based co-expression networks for 11 species (8 dicots, 2 monocots, and 1 ancestral species) with substantial anther RNA-seq coverage, using the conserved gene families defined above as nodes. Reproductive gene families were first grouped into five pathway categories—exine, intine, tapetum, pollen tube, and a broader “pollen development” group—based on *A. thaliana* annotations integrated from multiple databases (Fig. 4A; Fig. S6). For the exine, intine, pollen development, pollen tube, and tapetum pathways, GO annotations identified 66, 0, 264, 294, and 23 genes, respectively. Mercator identified 38, 5, 0, 19, and 5 genes, while PlantConnectome identified 52, 30, 241, 292, and 159 genes for the same pathways. In total, 98, 34, 444, 454, and 164 genes were assigned to each pathway. An edge between two families was drawn when the corresponding genes were co-expressed in at least two species, thereby retaining only conserved co-expression relationships. In the resulting network, we observed large numbers of non-reproductive or unannotated gene families (yellow nodes) that were strongly connected to reproduction-annotated families (colored nodes) across all pathways (Fig. 4A). Importantly, these co-expressed candidates were not restricted to a single clade: most were conserved across dicots, monocots, and ancestral lineages (D+M+A), as reflected both in their node shapes and in the strong enrichment of D+M+A families among non-reproductive nodes (Fig. 4B). This indicates that many uncharacterized, yet deeply conserved, gene families are tightly integrated into the core transcriptional programs of pollen development.

**Figure 4.**
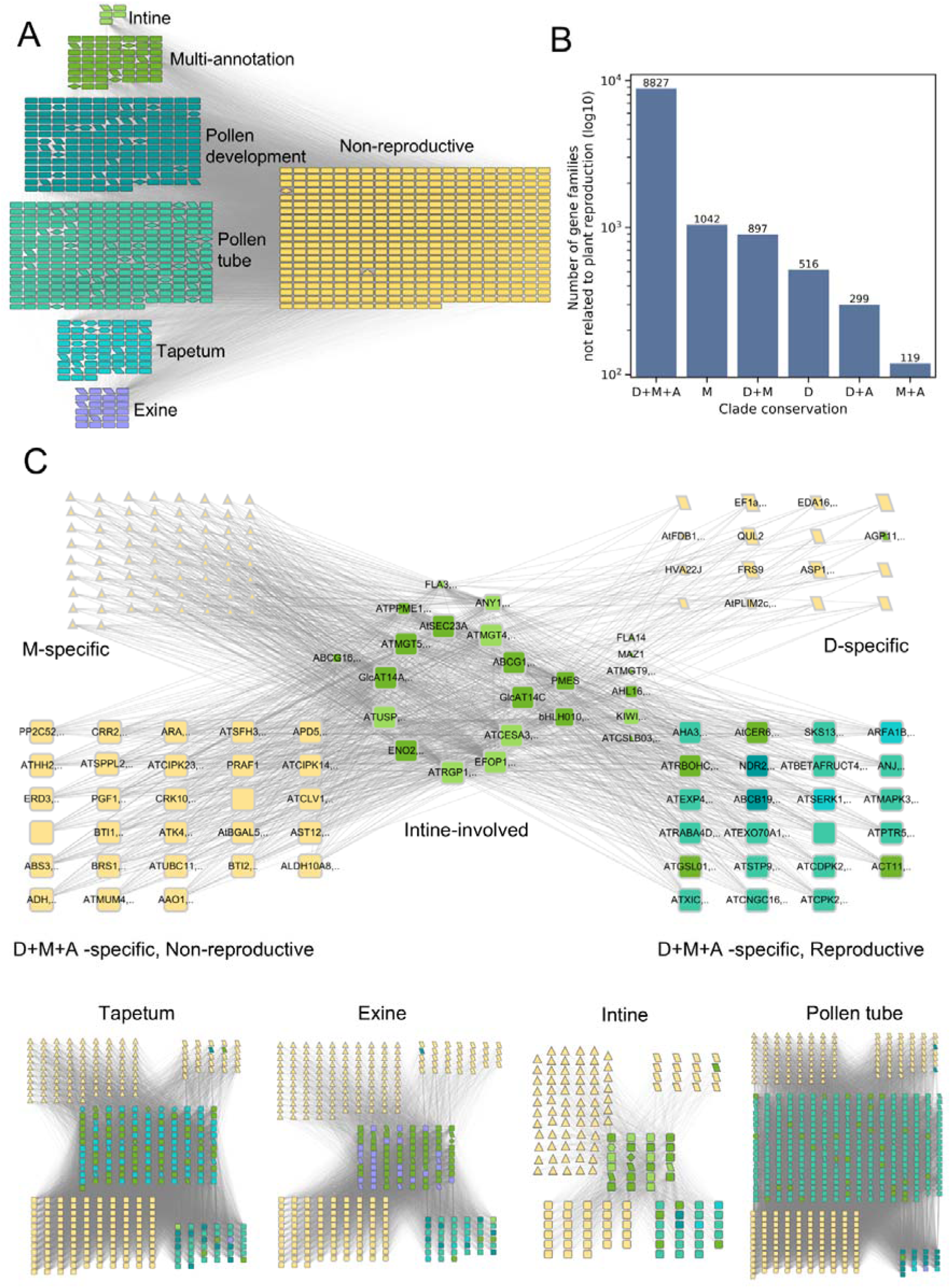
Identification of conserved pollen development gene co-expression networks (GCNs). (A) Non-reproductive or unannotated gene families (yellow nodes) co-expressed with reproduction-annotated gene families (green and purple nodes). From top to bottom, reproduction pathways correspond to intine, multi-annotation, pollen development, pollen tube, tapetum, and exine. Multi-annotation refers to gene families annotated with two or more terms among intine, pollen development, pollen tube, tapetum, and exine. Node shapes indicate different lineages, Dicots: parallelogram, Monocots: triangle, D+A: hexagon, D+M, diamond, D+M+A: rounded rectangle, M+A: V shape. (B) Number of non-reproductive or unannotated gene families conserved across major plant clades. (C) Conserved GCNs in intine-associated pathways. Node size represents the number of species in which the gene family is conserved.

To characterize pathway-specific modules in more detail, we visualized, for each pathway, the subset of monocot-specific, dicot-specific, fully conserved (D+M+A) non-reproductive gene families and previously annotated reproductive families that formed highly connected clusters (Fig. 4C; Fig. S7). Using the intine-associated network as an example (Fig. 4C), most pathway gene families showed high within-pathway conservation, as indicated by their large node sizes. We detected strongly conserved PME (pectin methylesterase) gene families that were co-expressed with pollen tube–related genes, consistent with their roles in cell wall remodeling during tube emergence and elongation.^29^ The network also highlighted coordinated expression of USP, RGP, and CESA gene families, which have been implicated in carbohydrate metabolism and cellulose synthesis,^5^ as well as gene families involved in calcium signaling (e.g., CIPK, CPK),^30,31^ vesicle trafficking (RAB/ARA),^32^ and additional wall-modifying enzymes (BGAL, PGF).^33,34^ These modules collectively support a conserved mechanistic model in which intine formation and pollen tube growth are controlled by tightly coupled regulation of signaling, trafficking, and cell wall biosynthesis genes.^5^

To probe the functional landscape of these co-expressed modules more systematically, we summarized the Gene Ontology (GO) annotations of gene families linked to each pathway using word clouds (Fig. S8). Each gene family was annotated based on experimentally supported or predicted GO terms; families lacking any annotation were designated as “unknown function.” Across pathways and lineages, we observed several conserved functional themes. In the D+M+A group, membrane-associated functions and ATP-binding activities were particularly prominent, consistent with the central roles of transporters, receptors, and ATP-dependent enzymes in pollen cell biology.^35–37^ By contrast, dicot-specific (D) and monocot-specific (M) families contained a relatively higher proportion of unknown functions (Fig. S8), suggesting that clade-restricted innovations in pollen development are still poorly characterized at the molecular level. Notably, monocot-specific candidates were enriched for ADP-binding and defense-related functions, hinting at lineage-specific rewiring of stress and signaling pathways in monocot pollen (Fig. S8). Together, these comparative co-expression analyses reveal deeply conserved transcriptional modules underpinning key aspects of pollen development, while simultaneously exposing lineage-restricted, largely uncharacterized gene families as promising targets for future functional studies (Fig. 4A–C; Fig. S7–S8).

### Divergent pollen traits and transcriptional programs in monocots and dicots

To place pollen-and anther-specific transcriptomes into an evolutionary and phenotypic context, we asked which pollen traits most consistently distinguish monocots and dicots (eudicots). We therefore performed a meta-analysis of pollen phenotype annotations from the PalDat database(www.paldat.org), focusing on structural and metabolic characteristics with sufficient sampling depth across clades (Fig. 5A). While all major lineages were largely consistent in producing monad pollen units, multiple traits showed significant clade-associated differences between monocots and dicots. For example, dicot pollen grains are generally smaller, with sizes predominantly distributed between 10–50 µm, whereas monocot pollen grains are typically larger, falling within the 50–100 µm range. In addition, dicots exhibited isopolarity patterns that were clearly distinct from those of monocots. Differences were also observed in aperture characteristics. Monocot pollen predominantly displayed a sulcate aperture type, while dicot pollen showed a higher number of apertures overall. When the pollen surface was examined using SEM, dicots more frequently exhibited perforate surface patterns, whereas monocots showed a higher prevalence of reticulate patterns. The Ubisch body, a precursor structure involved in sporopollenin deposition, was more frequently observed in monocots. In contrast, pollen kitt was more commonly detected in dicots. In the cytoplasmic region, dicot pollen contained higher starch levels and showed a slight increase in the occurrence of the tricellular (three-cell) structure. Some phenotypes did not show substantial differences between dicots and monocots, including the monad pollen unit, tectum, and intine structures.(Fig. 5A, Fig. S9)

**Figure 5.**
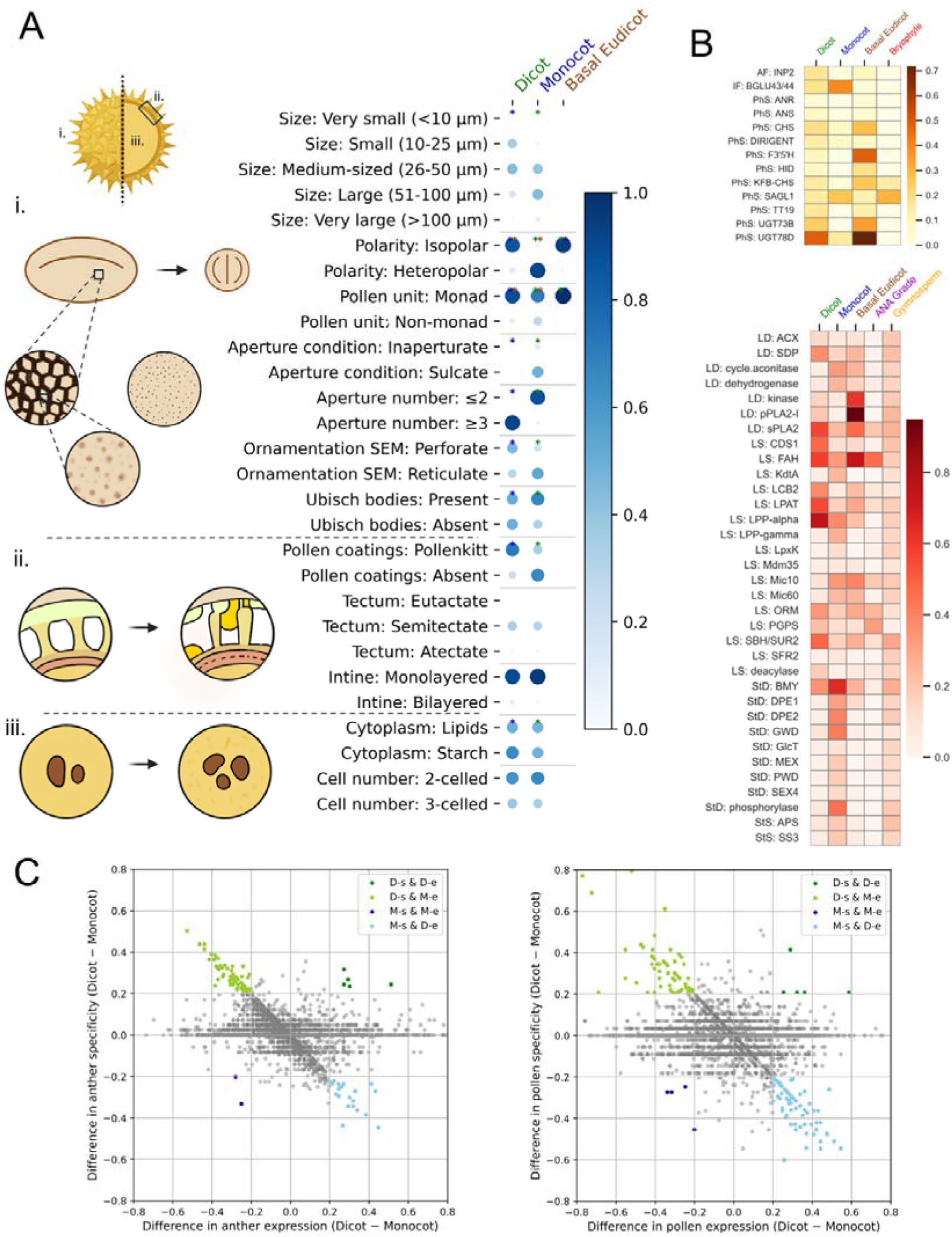
Evolutionary analysis of pollen-associated gene expression and structural traits. (A) Meta-analysis of pollen structural and metabolic phenotypic characteristics from the PalDat database. (B) Analysis of specifically expressed genes in anther (top, yellow) and pollen (bottom, red). Clade-specificity was calculated for dicots (eudicots) and monocots based on the phenotypic traits shown in panel B. Only genes with clade specificity ≥ 0.6 and statistically significant differences (p < 0.05, Mann–Whitney U test) were retained. AF – Aperture Formation, if-Intine Formation, PhS – Phenolics Synthesis, LD - Lipid Degradation, LS – Lipid Synthesis, StS – Starch Synthesis, StD-Starch Degradation (C) Identification of Dicot-and Monocot-specific genes in anther (left) and pollen (right). D-s and M-s denote gene functions specific to dicots and monocots (mean SPM ≥ 0.8), respectively, whereas D-e and M-e denote gene functions expressed in dicots and monocots (0 < mean SPM < 0.8), respectively.

We next tested whether the lineage-level trait differences inferred from PalDat are reflected in lineage-biased expression of corresponding biological pathways. To do this, we manually curated sets of genes linked to the trait-associated processes (e.g., aperture formation, intine formation, phenolics metabolism, lipid synthesis/degradation, and starch synthesis/degradation) using Mercator annotations, and compared their tissue-specific expression patterns between monocots and dicots. In anther-specific gene sets, dicots showed elevated expression of genes INP2, ANR, ANS, CHS, DIRIGENT, F3’5’H, HID, KFB-CHS, TT19, UGT73B, and UGT78D associated with aperture formation and phenolics synthesis (Fig. 5B, top),^38–48^ consistent with the strong clade differentiation in aperture-related traits and wall-associated chemistry observed in the phenotypic meta-analysis (Fig. 5A). In pollen-specific gene sets, monocots showed increased expression of starch degradation-associated genes including BMY, DPE1, DPE2, GWD, GlcT, MEX, PWD, SEX4, APS, SS3, and cytosolic alpha-glucan phosphorylase (Fig. 5B, bottom, Fig. S10 for individual genes),^49–55^ aligning with the PalDat-derived differences in cytoplasmic content and inferred metabolic profiles (Fig. 5A).

Finally, to resolve whether dicot–monocot differences primarily reflect shifts in overall expression versus shifts in strict organ specificity, we compared clade bias along two axes: differential expression (0 < median SPM < 0.8) and differential specificity (median SPM ≥ 0.8) for each gene function category, as defined by MapMan (Table S6, S7 shows the results for pollen and anthers, respectively). Functions near x = 0 and y = 0 represent broadly conserved deployment between dicots and monocots in both expression and specificity (Fig. 5C). Along the axes, positive values indicate dicot-enriched expression (x) or specificity (y), whereas negative values indicate monocot enrichment and specificity (Fig. 5C). Notably, they also reveal a diagonal pattern spanning the second and fourth quadrants, corresponding to functions that are strongly specific in one clade yet detectably expressed in the other (light green and light blue dots)—consistent with partial conservation of transcriptional deployment but divergence in regulatory restriction. This framework highlighted multiple gene functions that are both highly expressed and highly specific in either monocots or dicots, representing candidate pathways most likely to underpin lineage-distinct pollen traits (Fig. 5C; Table S6,S7). Specifically in pollen, ABCG31, bHLH class-Ib transcription factor, SBT4, AP180, NLR, and SRC2-clade calcium sensor genes showed strong dicot enrichment, appearing in the upper right region of the plot (dark green dots). These genes are associated with abscisic acid transport, transcriptional regulation, protein turnover, vesicle trafficking, immune receptors, and calcium-dependent signaling.^56–61^ In contrast, CYP94C, class-C endo-1,4-beta-glucanase, FUT, and LAV-ABI3 transcription factor genes showed monocot enrichment (dark blue dots), with functions related to jasmonic acid metabolism, cell wall modification, xyloglucan biosynthesis, and transcriptional regulation.^62–65^ A similar pattern was observed in anther, where ADS/FAD5, UGT78D, SBT4, AP180, and DMP8-9 genes showed dicot enrichment, functioning in fatty acid desaturation, flavonol biosynthesis, proteolysis, vesicle trafficking, and gamete fusion.^48,58,59,66,67^ In contrast, LAV-ABI3 transcription factor and GAPLESS genes showed monocot enrichment, associated with transcriptional regulation and plasma membrane tethering during Casparian strip formation.^62,68^

### Experimental validation of the conserved anther and pollen genes

To test whether our pipeline enriches for genes required for pollen development and function, we characterized loss-of-function mutants for 20 genes prioritized from the conserved pollen/anther-specific families. Conservation patterns of these genes and their families are summarized in Table S8. We deliberately selected families spanning diverse conservation profiles—ranging from fully conserved (D+M+A) to dicot-specific and dicot+monocot–specific—while varying family size and the number of species in which the family was expressed or specific in anther or pollen. Phenotypic assays focused on key readouts of male reproductive performance: pollen viability (Alexander staining), pollen germination rate, pollen tube length in vitro, and pollen morphology by SEM (Fig. S11–S12). In addition, we assessed whole-plant fertility via silique length, seed number, and seed abortion rate (Fig. 6A,D,E; Fig. S12).

**Figure 6.**
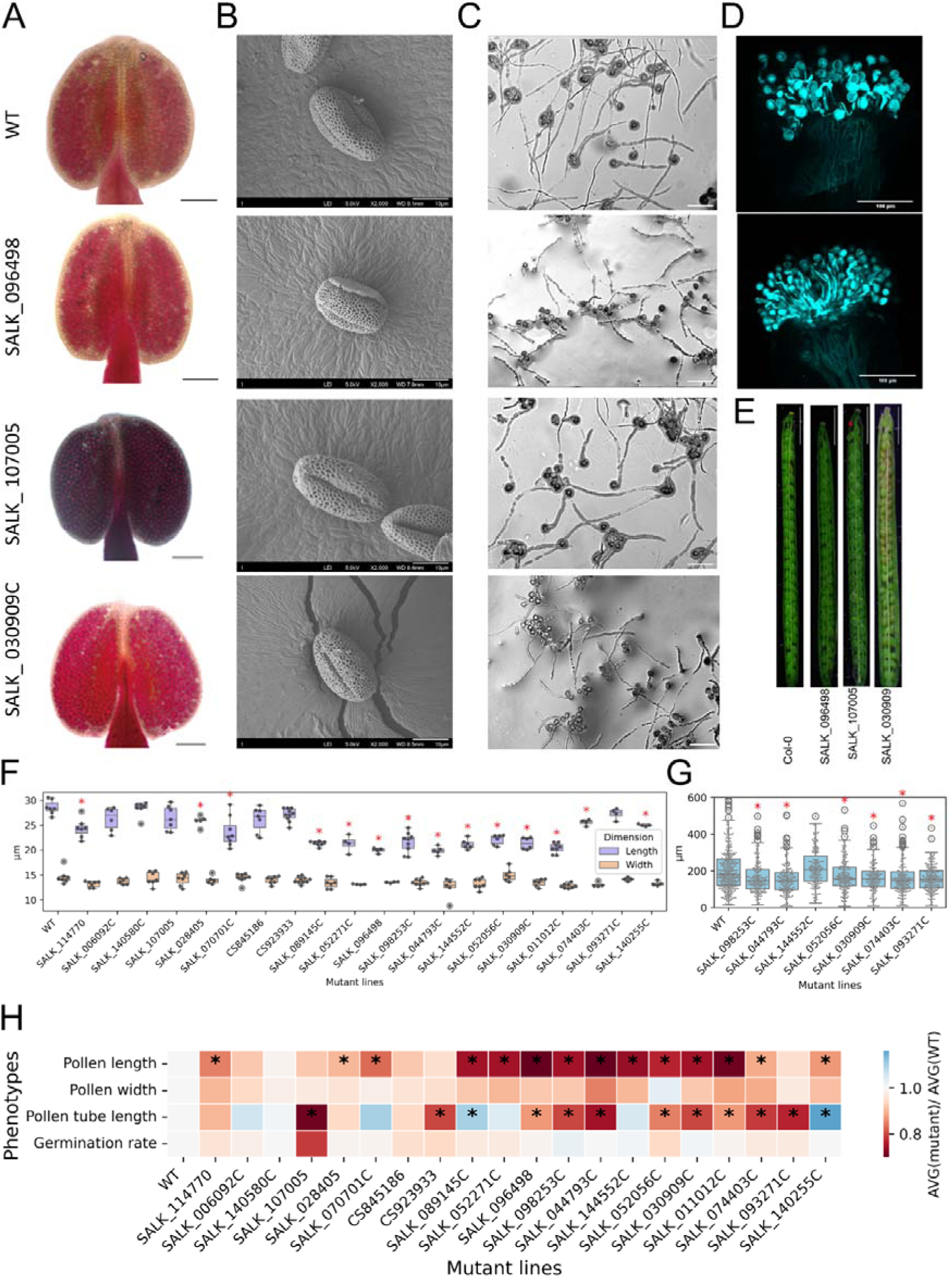
Different experimental characterization of 20 homozygous mutant lines. The experimental characterization includes: (A) pollen viability, (B) pollen scanning electron microscopy (SEM) imaging, (C) pollen tube length, (D) stigma pollination rate and (E) silique morphology analysis. (F) Pollen length and width and (G) pollen tube length statistical analysis, where each point represents one pollen/pollen tube. The red asterisk indicates lines significantly (BH-corrected p < 0.05, Welch’s t-test) shorter than the wild type control. (H) shows a summary of phenotypes observed in 20 homozygous mutant lines, where each cell represents the ratio of the mutant average, divided by wild type average. The asterisk indicate significantly (BH-corrected p < 0.05, Welch’s t-test) different measurements in the mutant vs. the wildtype.

Across the 20 lines, we did not observe strong defects in pollen viability, stigma pollination rate, seed abortion, or seed germination (Fig. 6A,D,E; Fig. S12), suggesting that most tested genes are not individually essential for gross male fertility under our growth conditions, consistent with substantial redundancy in many conserved families. In contrast, several mutants showed clear quantitative defects in pollen morphology and tube growth: we detected significant reductions in pollen grain length (Fig. 6F) and pollen tube length (Fig. 6G) in multiple lines, with a summary of all phenotypic outcomes provided in Fig. 6H. For example, the SALK_096498 line, carrying a T-DNA insertion in AT5G60510 (Table 1), displayed normal pollen viability and stigma pollination (Fig. 6A,D) but produced shorter pollen grains and shorter pollen tubes (Fig. 6B,C). These results support the prioritization strategy by linking conserved pollen/anther-specific genes to measurable defects in pollen morphology and tube growth, while also highlighting the prevalence of subtle, quantitative phenotypes rather than complete loss of fertility for many individual mutants.

**Table 1.** AGI identifiers, T-DNA insertion lines, gene descriptions in TAIR and phenotypic summary of 20 mutant lines.

| AGI identifier | T-DNA insertion lines | TAIR10 gene description | Phenotype description |
| --- | --- | --- | --- |
| AT1G18960 | SALK_074403<br>C | myb-like HTH transcriptional regulator family protein;(source:Araport11) | Decreased pollen and pollen tube length |
| AT1G75030 | SALK_089145<br>C | encodes a PR5-like protein | Decreased pollen and increased pollen tube length |
| AT3G51590 | SALK_052271<br>C | Encodes a member of the lipid transfer protein family. Proteins of this family are generally small (~9 kD), basic, expressed abundantly and contain eight Cys residues. The proteins can bind fatty acids and acylCoA esters and can transfer several different phospholipids. They are localized to the cell wall. The LTP12 promoter is active exclusively in the tapetum during the uninucleate microspore and bicellular pollen stages. | Decreased pollen length |
| AT3G57620 | SALK_093271<br>C | Galactose oxydase; may function in tissues that require mechanical reinforcements in the absence of lignification. | Decreased pollen tube length |
| AT5G60510 | SALK_096498 | Undecaprenyl pyrophosphate synthetase family protein;(source:Araport11) | Decreased pollen and pollen tube length |
| AT3G54800 | SALK_140255<br>C | Pleckstrin homology (PH) and lipid-binding START domains-containing protein;(source:Araport11) | Decreased pollen and increased pollen tube length |
| AT5G49920 | SALK_098253<br>C | Octicosapeptide/Phox/Bem1p family protein;(source:Araport11) | Decreased pollen and pollen tube length |
| AT3G56180 | SALK_044793<br>C | LURP-one-like protein (DUF567);(source:Araport11) | Decreased pollen and pollen tube length |
| AT4G27790 | SALK_144552<br>C | Calcium-binding EF hand family protein;(source:Araport11) | Decreased pollen length |
| AT2G35215 | SALK_052056<br>C | plant/protein;(source:Araport11) | Decreased pollen and pollen tube length |
| AT3G23350 | SALK_030909<br>C | ENTH/VHS family protein;(source:Araport11) | Decreased pollen and pollen tube length |
| AT2G03840 | SALK_011012<br>C | TET13 encodes a member of the TETRASPANIN gene family that is expressed in the hypophysis, QC, root stem cells, lateral root primordia and is involved in primary root growth and lateral root development. | Decreased pollen and pollen tube length |
| AT5G25530 | CS923933 | DNAJ heat shock family protein;(source:Araport11) | Decreased pollen tube length |
| AT5G09430 | SALK_114770 | alpha/beta-Hydrolases superfamily protein;(source:Araport11) | Decreased pollen length |
| AT2G27180 | SALK_028405 | hypothetical protein;(source:Araport11) | Decreased pollen length |
| AT3G06778 | SALK_006092<br>C | Chaperone DnaJ-domain superfamily protein;(source:Araport11) |  |
| AT5G02390 | SALK_140580<br>C | Target promoter of the male germline-specific transcription factor DUO1. |  |
| AT4G10850 | CS845186 | Nodulin MtN3 family protein |  |
| AT5G41890 | SALK_070701C | GDSL-motif esterase/acyltransferase/lipase. Enzyme group with broad substrate specificity that may catalyze acyltransfer or hydrolase reactions with lipid and non-lipid substrates. | Decreased pollen length |
| AT4G38230 | SALK_107005 | member of Calcium Dependent Protein Kinase | Decreased pollen tube length |

## Discussion

Pollen transcriptomes are strongly enriched for genes of unknown function across species^8^, indicating that even the conserved core of male reproduction remains incompletely understood. Motivated by the essential role of pollen in plant fertility and emerging uses of pollen as a renewable material, we built a cross-species atlas of 90 land plants, combining public and newly generated datasets. Within this phylogenetically broad framework, pollen and anther consistently emerged as “specialized transcriptomes”: across species, they rank among the organs with the highest fractions of tissue-specific genes (Fig. 2C), underscoring the unusually tight transcriptional restriction that underpins male reproduction.

Organ specificity proved to be a reliable large-scale proxy for function. Leaf-specific genes were enriched for canonical photosynthesis pathways, whereas pollen-and anther-specific genes were enriched for reproductive processes (Fig. 2D), showing that PO-based organ harmonization plus SPM recovers biologically meaningful patterns despite heterogeneous source data. Within this space, we identified highly conserved pollen/anther-specific gene families, many still uncharacterized (Fig. 3A), highlighting a substantial “known-unknown” component in core male reproductive programs. Functional partitioning of conserved families matched tissue division of labor: anther-specific families were enriched for developmental and structural processes (e.g. wall and exine formation), whereas pollen-specific families were enriched for germination, tube growth, and guidance (Fig. 3B).

Our comparative co-expression analysis revealed that these conserved families are embedded in deeply conserved transcriptional modules. Anther-based GCNs across 11 species identified many non-reproductive or unannotated families that are strongly co-expressed with exine, intine, tapetum, pollen tube, and general pollen development genes (Fig. 4A–C). Most of these candidates are conserved across dicots, monocots, and early-diverging lineages, indicating that core pollen modules integrate both well-annotated and unknown components. At the same time, we observed clear monocot–dicot divergence: PalDat trait differences in aperture patterns, wall organization, and cytoplasmic reserves (Fig. 5A) were mirrored by clade-biased specificity and expression in corresponding gene sets (Fig. 5B,C), suggesting that shifts in pollen form have been accompanied by both rewiring of shared gene families and the recruitment of lineage-restricted families.

Loss-of-function analysis in *A. thaliana* broadly supported our prioritization strategy. Many mutants showed subtle or no effects on pollen viability or fertility, consistent with gene-family redundancy and context-dependent roles, whereas recurrent defects in pollen morphology and tube growth (Fig. 6; Fig. S11-12) reinforced that conserved pollen-specific genes disproportionately impact tube-related phenotypes. While it is difficult to explain phenotypes with a single parameter of conservation, we observed that gene families showing reduced pollen tube length tended to have larger family sizes and showed high conservation across both pollen and anther in terms of expression and specificity levels. In contrast, gene families linked to reduced pollen length were relatively smaller in size and displayed more dicot-preferred or incompletely conserved patterns in pollen or anther. Gene families associated with the mixed phenotype exhibited a broader range of conservation levels, along with greater variation in gene family size (Table S8).

Our study has limitations. The RNA-seq compendium is heterogeneous in genotype, environment, and developmental staging; “anther” and “pollen” samples differ in purity and cell-type composition; and the stringent SPM ≥ 0.8 threshold is necessarily heuristic. Mutant validation relied mainly on single T-DNA alleles without complementation. However, the breadth of our sampling (90 diverse species) mitigates many dataset-level artefacts: false positives are unlikely to recur as tissue-specific across many species, whereas truly conserved organ-specific families are repeatedly recovered. Overall, the expression atlas, organ-specific gene sets, conservation scores, and co-expression modules provide a reusable framework and shortlist resource for dissecting male fertility, informing hybrid breeding and stress-resilience strategies, and guiding mechanistic studies of pollen as both a biological and engineering material.

## Materials and methods

### RNA extraction and in-house transcriptome assembly

Our RNA-seq data of 90 different species consists of 8 in-house sequencing data and 82 publicly available databases (NCBI Sequence Read Archive)^69^ and Plant Expression Omnibus^70^.

We sampled 5 dicot and 3 monocot species representing eight families from the Singapore Botanic Gardens. Collected organs included anthers and other representative tissues such as leaf, root, stem, seed, and fruit (Table S2). Plant organs were first dissected and immediately frozen in liquid nitrogen. The samples were then stored in 15-ml Falcon tubes at −80 °C to prevent RNA degradation. Total RNA was extracted from tissues ground into a fine powder under liquid nitrogen. RNA quality assessment and library construction were performed by Novogene (Singapore).

The de novo transcriptome assembly pipeline for generating pseudo-genomes of the 8 Singaporean local plants can be found in Fig S2. In the pipeline, raw reads were first quality-filtered using FastP to remove low-quality bases, adaptor sequences, and erroneous reads. For each organ, assemblies were generated independently using both SOAPdenovo-Trans (v.1.04) and Trinity (v2.15.1), and redundant transcripts were reduced with the EvidentialGene pipeline (trformat.pl and tr2aacds.pl) to retain the most complete coding sequences. Assembly quality was evaluated with BUSCO (v.5.8.0). Transcripts from all organs were then concatenated, and tr2aacds.pl was applied again at the species level to obtain the final transcript set. Transcript abundance was estimated with Kallisto (v.0.51.0), and lowly expressed transcripts (average TPM < 1 per organ) were removed. Coding sequences with GC content below 40% or above 60% were excluded, and final decontamination was performed using BLAST (v.2.16.0+) against the NCBI Core Nucleotide Database, removing sequences matching non-viridiplantae species (identity > 70%, e-value < 1e-10).

### Public RNA-seq data collection, processing, and sample annotation

For publicly available RNA-seq data, all selected species contained at least one sample annotated with male reproductive organs directly related to pollen development, including anther, pollen, microspore, and pollen tube. RNA-seq data from other major organs, including leaf, stem, root, seed, fruit, and floral tissues, were additionally collected to construct SPM matrices. When a sample lacked direct annotation to a major organ, its assignment was inferred from descendant Plant Ontology (PO) terms through “part_of” or “is_a” relationships.

To quantify transcript expression level of the RNA-seq data, Kallisto and LsTrAP-Cloud were used to generate Transcript per million (TPM) values. The quality control of the TPM matrices were done by using the pipeline in Fig S3. Samples with the number of reads < 2,000,000 and <15 to 30 mapping percentage were removed.

### Identification of organ-specific and expressed genes

The TPM matrices were used to generate specificity measures (SPM) to identify genes specific to different organs. For the genes with TPM>2, the SPM was defined as:

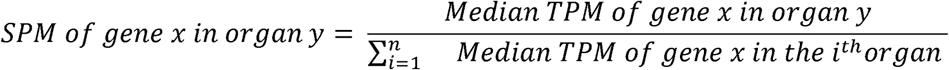

where n denotes the number of different tissues in a species and y denotes a tissue of interest.^8^ SPM ranges from 0 (ubiquitous expression) to 1 (organ-specific expression). In our analysis, genes with SPM larger than 0.8 were defined as organ-specific genes and genes with SPM smaller than 0.8 and larger than 0 were defined as organ-expressed genes.

### Identification of pollen-and anther-specific gene families

MMseqs2 (v.18), a fast and sensitive protein sequence searching tool,^71^ was used with default settings to identify gene families from the proteomes of the 90 species. The pollen-or anther-specific gene families were identified using tissue-expressed genes (0.8 > SPM > 0) and tissue-specific genes (SPM ≥ 0.8). The degree of conservation was identified with the number species with pollen-or anther-specific genes. For instance, if a pollen-specific gene family’s degree of conservation is 10, there are 10 different species which had at least one pollen-specific gene belonging to the gene family. Complete information of the pollen-and anther-specific gene families can be found in Table S4 and S5.

### Pathway enrichment analysis of tissue-specific genes

Mercator4 (v6.0) was used to identify pathway enrichment of different organ-specific genes.^72^ For every species, 30% of genes with high organ-specificity (SPM ≥ 0.8) from each major organ were randomly sampled five times. Thereafter, cross-species conservation at the second-level MapMan bins were calculated. The cross-species conservation for each organ i was defined by the number of species with at least one organ i specific gene divided by the total number of species with the organ i RNA-seq data in each corresponding second-level MapMan bin. These processes were repeated 5 times, with mean and standard deviation across replicates shown as dots and error bars, respectively. A two-sample permutation test (10,000 permutations) was applied to evaluate differences in mean values between the target genes and non-target genes. In the photosynthesis-related bins, target genes were leaf-specific genes and non-target genes were all the other tissue-specific genes. In the case of reproduction-related bins, permutation tests were performed between pollen/anther-specific genes and non-reproductive organs (for “pollination and fertilization” and “gametogenesis”), and between seed-specific and non-seed genes for the remaining reproductive terms.

The pathway enrichment for the conserved pollen-and anther-specific genes was calculated using experimentally verified *A. thaliana* GO Slim terms^73^. First, the conservation degree (number of different pollen (anther)-specific genes within a gene family) was identified for all the pollen (anther)-specific *A. thaliana* genes with any GO annotations. Each GO annotation was assigned with conservation degree summation of corresponding genes and plotted into a wordcloud figure.

### Construction and analysis of gene co-expression networks

To study pollen and anther genes at a network scale, we generated gene co-expression network (GCN) using TEA-GCN^74^, with anther RNA-Seq data of 11 species. *A. thaliana* GO Slim^73^, Mercator4 (v6.0)^72^, and PlantConnectome^75^ were used to identify genes within different pollen-related pathways including exine, intine, pollen development, pollen tube, and tapetum. In total, 98, 34, 444, 454, and 164 genes were identified in each pathway.

Gene families, including those containing pathway genes (pathway gene families), were identified based on MMseqs2 (v.18) clustering results.^71^ Nodes represent gene families that are co-expressed with pathway gene families, while edges indicate co-expression between two different gene families observed in at least two different species. The co-expression cut-off was set at a zScore (Co-exp_Str_MR) ≥ 1.7. GO term annotations for each node were inferred using InterProScan (v.5.76-107.0) based on the representative sequence of the family.^76^

### Evolutionary analysis of gene functions and pollen phenotypes

Pollen phenotype datasets of 4548 species were collected from PalDat (www.paldat.org) with our in-house script. For each feature, clades with ≥30 data points were retained. From each clade, 50% of the minimum dataset size was randomly sampled 100 times, with counts normalized to the sampling size. Contingency tables were evaluated using Chi-square tests (≥3 categories) and Fisher’s exact tests (2 categories). *p < 0.05.

Dicot-and monocot-specific or expressed genes were identified using the maximum SPM value for each MapMan bin (Tables S6 and S7). Within each clade, gene function importance was classified into four levels: 1, missing annotation; 2, absent or low expression (TPM < 2); 3, expression without specificity (0 < SPM < 0.8); and 4, tissue-specific expression (SPM ≥ 0.8). Differences between monocot and dicot specific or expressed functions were calculated by subtracting the level 3 and level 4 values between the two clades.

### Growth and genotyping of *A. thaliana* mutants

*A. thaliana* plants were grown in the greenhouse under a light cycle of 16 h light and 8 h dark at 22 °C. *A. thaliana* ecotype Columbia-0 (Col-0) was taken as the wild type (WT). Prior to observing phenotypes of mutants, genotyping was conducted to only select homozygous lines. First, genomic DNAs were extracted from young leaves of each plant. Subsequently, PCR was conducted to WTs and T-DNA inserted mutant genes. Finally, the existence of the amplified genes in WTs, heterozygous, and homozygous lines were identified through gel electrophoresis. The list of primer sequences for identifying homozygous lines are in Table S9.

### Morphological analysis

Pollen grains were collected using an in-house pollen collector after the flowering stage. Three different-sized filters: 60, 35, and 10 μm-sized filters were used to capture pollen grains. To preserve original morphology of native or defatted pollen samples, laboratory-grade cotton swabs were used to gently transfer the samples onto carbon adhesive tape mounted on aluminum stubs. Subsequently, the mounted samples were coated with a thin layer of platinum using a sputter coater for 90 seconds at a current of 20 mA. The prepared samples were then examined under a scanning electron microscope (JSM-6700F, JEOL, Japan) for morphological analysis.

### Pollen germination

Mature pollen grains from newly opened flowers of 5-to 6-week-old *A. thaliana* were cultured on a solid pollen germination medium containing 10% (w/v) sucrose, 0.5 mM calcium chloride (CaClC), 0.5 mM potassium chloride (KCl), 0.1 mM magnesium sulfate (MgSOC), 0.01% boric acid (H_3_BO_3_), and 1.5% agar (pH 7.5) and germinated in a incubator at 28°C.

### Alexander staining for pollen viability assessment

Pollen grains and anthers from *A. thaliana* flowers were immersed in 70% ethanol for 1 minute. After removing ethanol with absorbent paper, the samples were incubated in Alexander staining solution supplemented with 0.5% (v/v) Triton X-100 and kept in the dark to complete the staining process.

### Aniline blue staining for stigma pollination rate measurement

Stigmas from pollinated flowers were fixed in a solution of anhydrous ethanol and glacial acetic acid (3:1 ratio) for 2 hours. After fixation, the samples were sequentially washed with a graded ethanol series (75%, 50%, 30%) for 10 minutes each, followed by rinsing with sterile water for 10 minutes. The stigmas were then softened in 8 M sodium hydroxide (NaOH) for 12 hours, rinsed thoroughly with water, and stained with 0.1% aniline blue solution.

### Fertility Analysis

Siliques collected from 7-to 8-week-old *A. thaliana* plants were dissected under a stereo microscope using fine forceps and imaged.

## Acknowledgments

We acknowledge Novo Nordisk Foundation Starting grant to M.M., Singaporean Ministry of Education Tier 3 grant to J.M., E.K., N.J.C. We thank Singaporean Botanic Garden for their help with plant sampling. We thank Q.W.Tan for providing the framework for transcriptomic assembly.

## Data availability

The raw sequencing data are available at E-MTAB-, and the coding sequences, expression matrices, and SPM matrices are available via Figshare at doi: 10.6084/m9.figshare.31333153

## Author contributions

E.K., R.S.S., J.Y.T., B.C.H., and L.C. were involved in the sampling of plants. J.M. was involved in the bioinformatical data collection and analysis. J.M. was involved in genotyping of mutants. E.K., Z.S., and Y.Z. were involved in characterization of mutants. W.P., S.D., Z.X., T.C.C.K., and J.M. assisted characterization of mutants. T.J.H. performed SEM imaging of mutants. T.C.C.K assisted collection of GCN data. J.M. and M.M. wrote the paper with help from N.J.C., M.M. conceptualized and supervised the project.

## Supplementary tables

Table S1. Organ, SRA run, and phylogenetic summary of RNA-seq data from 82 species collected from public databases

Table S2. Organ and phylogenetic summary of RNA-seq data from 8 local species collected in Singapore

Table S3. BUSCO summary statistics for the de novo transcriptome assemblies of the 8 Singaporean local species.

Table S4. Conservation information of gene families showing pollen expression (0 < SPM < 0.8) or pollen specificity (SPM ≥ 0.8)

Table S5. Conservation information of gene families showing anther expression (0 < SPM < 0.8) or anther specificity (SPM ≥ 0.8)

Table S6. MapMan functional annotations of genes with maximum pollen SPM across 44 species with available pollen RNA-seq data

Table S7. MapMan functional annotations of genes with maximum anther SPM across 58 species with available anther RNA-seq data

Table S8. Cross-species conservation status of the gene families associated with the 20 mutant lines

Table S9. Primer sequences obtained from the Salk SIGnAL database for identifying homozygous lines.

## Supplemental data

1. **Zip file with all CDS files of the 8 species**
2. **Zip file with the exp mats of the 8 species**
3. **SPM for all genes**

## Supplementary figures

**Figure S1.**
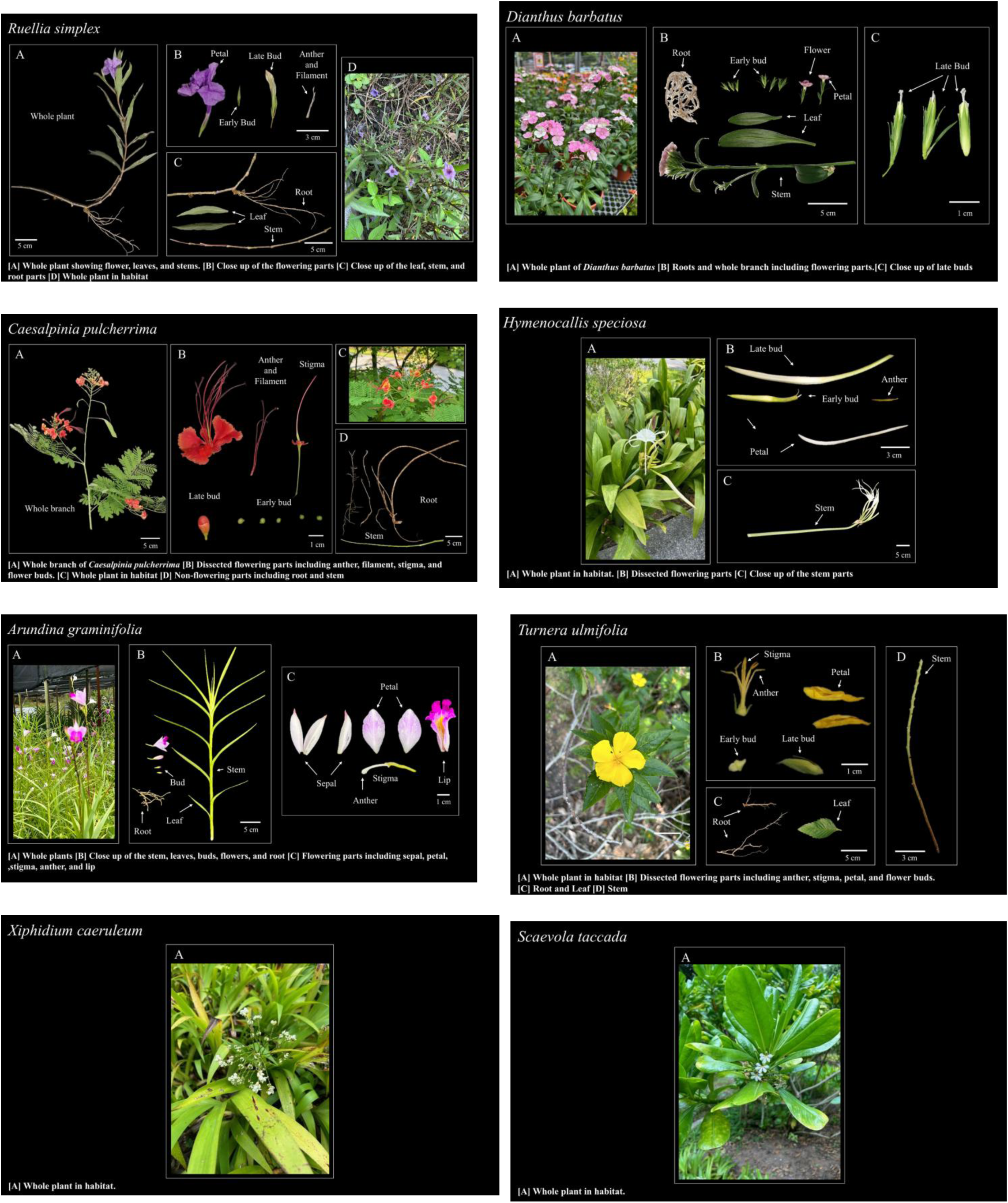
Representative images of the eight Singaporean local plant species and the organs used for RNA sequencing. Samples were collected from the Singapore Botanic Gardens. The in-house collection includes five dicot species (*Dianthus barbatus*, *Caesalpinia pulcherrima*, *Ruellia simplex*, *Turnera ulmifolia*, and *Scaevola taccada*) and three monocot species (*Arundina graminifolia*, *Xiphidium caeruleum*, and *Hymenocallis speciosa*).

**Figure S2.**
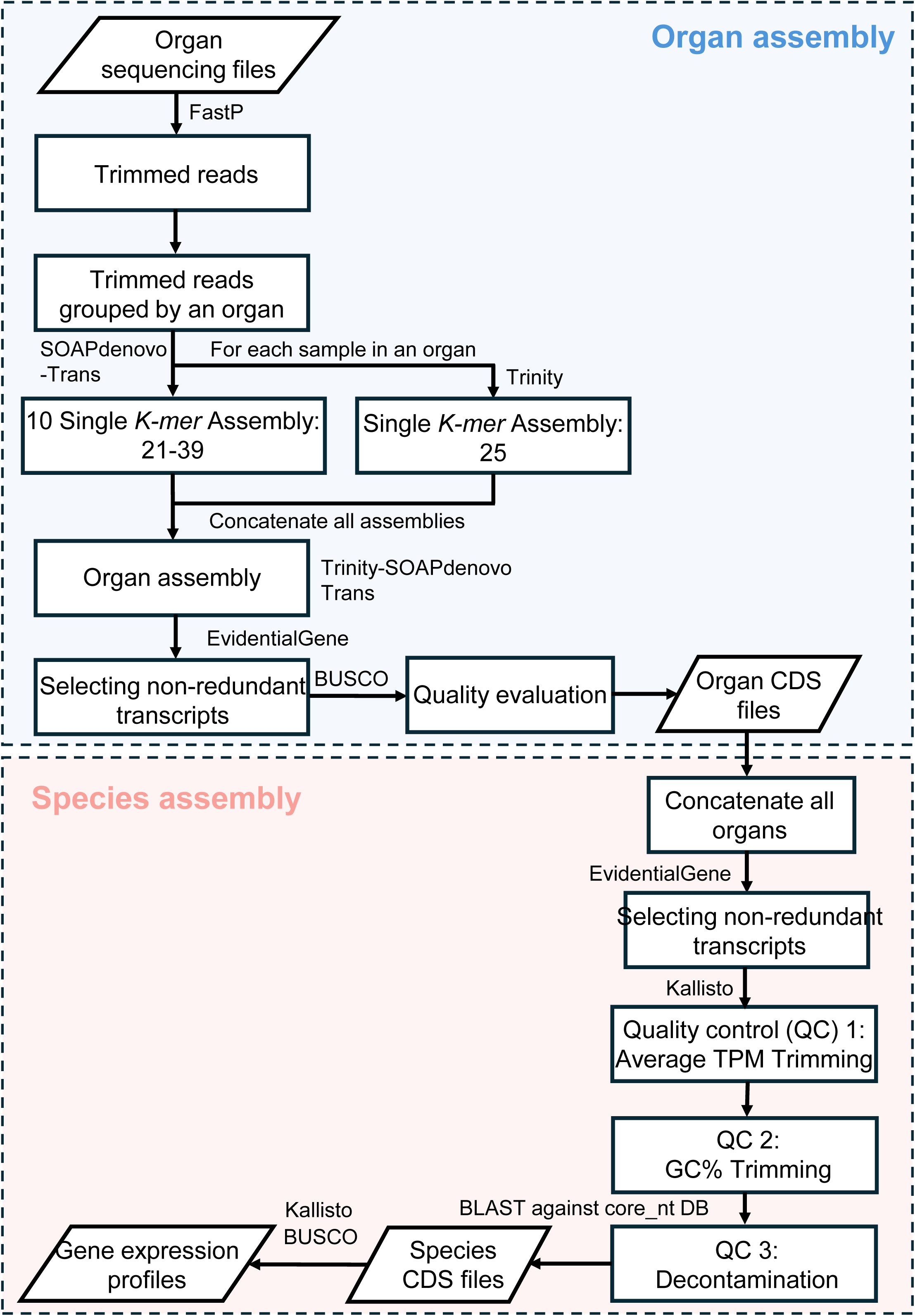
Flowchart of the de novo transcriptome assembly pipeline

**Figure S3.**
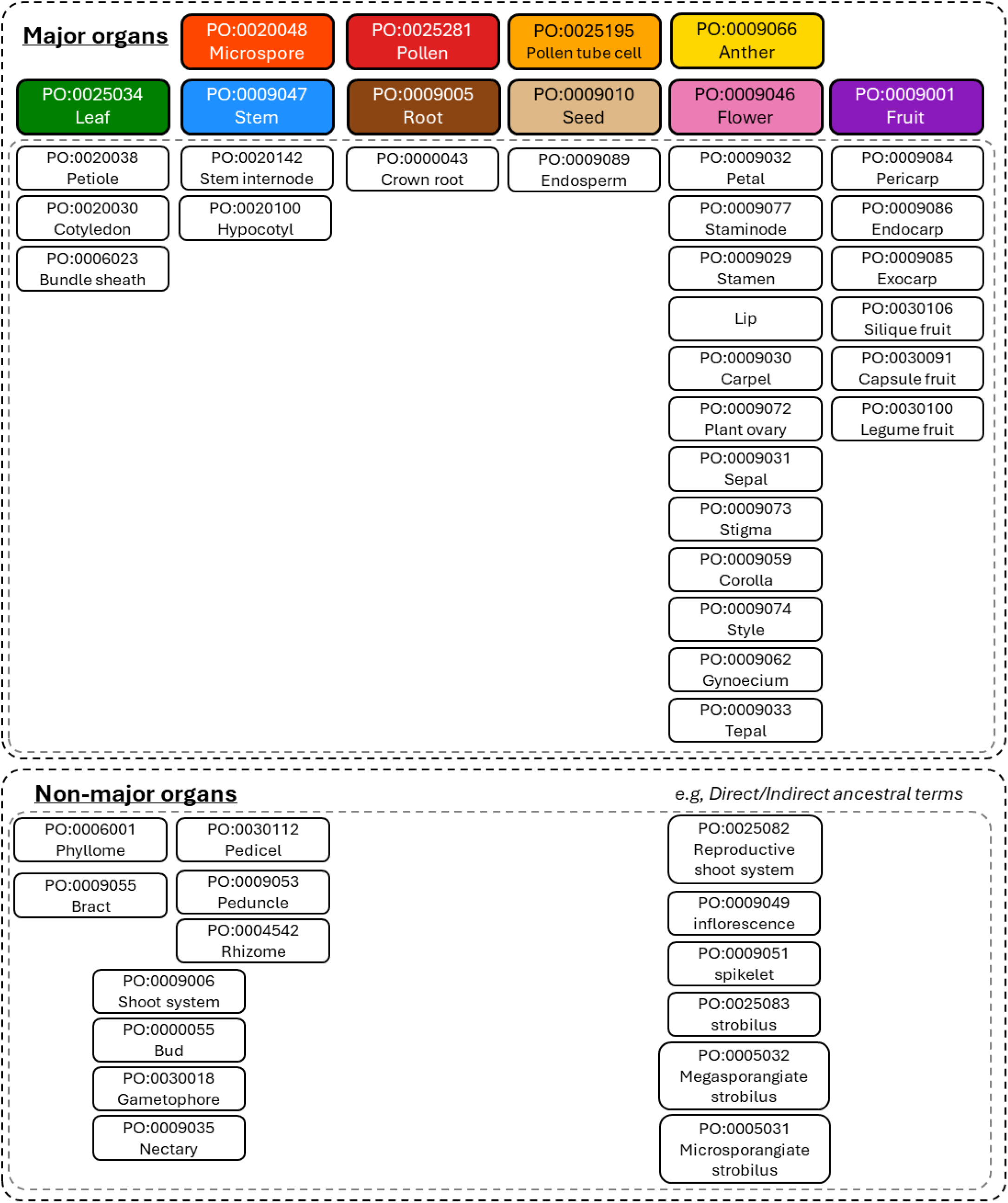
Classification of plant organs used for SPM calculation. Plant organs were categorized based on Plant Ontology (PO) terms and grouped into ten major organ types: microspore, pollen, pollen tube cell, anther, leaf, stem, root, flower, seed, and fruit. All descendant terms associated with each major organ (“part_of” or “is_a”) were recursively aggregated under the corresponding major category. Terms not falling within these categories, including direct or indirect ancestral terms (representing temporally earlier developmental stages) and spatially or temporally too broad/specific descriptors, were considered non-major organs. To ensure consistent comparisons of organ-specific genes across species, the number of distinct organs assigned to each species was limited to a maximum of ten. List of all non-major organs can be found from Table S1

**Figure S4.**
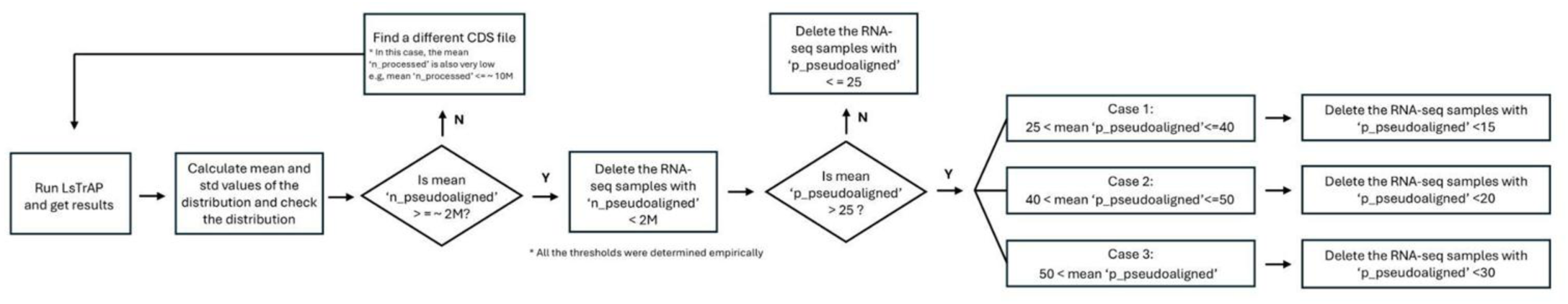
Quality control pipeline for gene expression analysis using Kallisto RNA-seq samples were evaluated using LSTrAP based on pseudoalignment metrics. Samples with low numbers of pseudoaligned reads(n_pseudoaligned) or low pseudoalignment rates(p_pseudoaligned) were removed using empirically determined thresholds. If overall alignment quality was insufficient, an alternative CDS reference was selected and the process was repeated.

**Figure S5.**
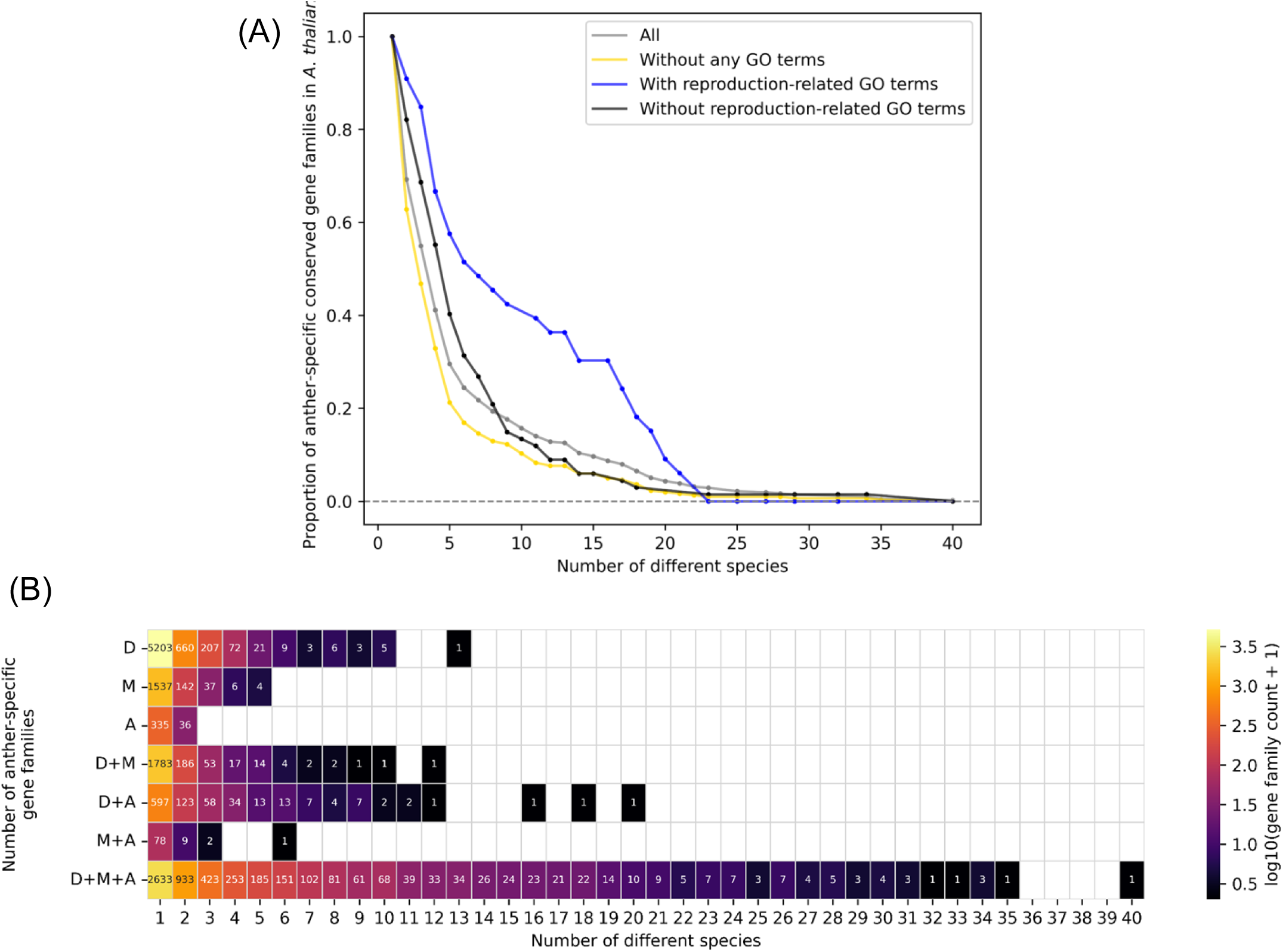
Conservation of anther-specific gene families in (A) *A. thaliana* and (B) all species with available anther RNA-seq data (A) Proportion of total and experimentally unstudied conserved anther-specific gene families (SPM ≥ 0.8) in *Arabidopsis thaliana*. The number of different species denotes the number of species containing at least one anther-specific gene within a clustered anther-specific gene family. (B) Number of anther-specific gene families conserved across major plant clades: dicots, monocots, and ancestral lineages (including bryophytes, ANA-grade, angiosperms, and basal eudicots).

**Figure S6.**
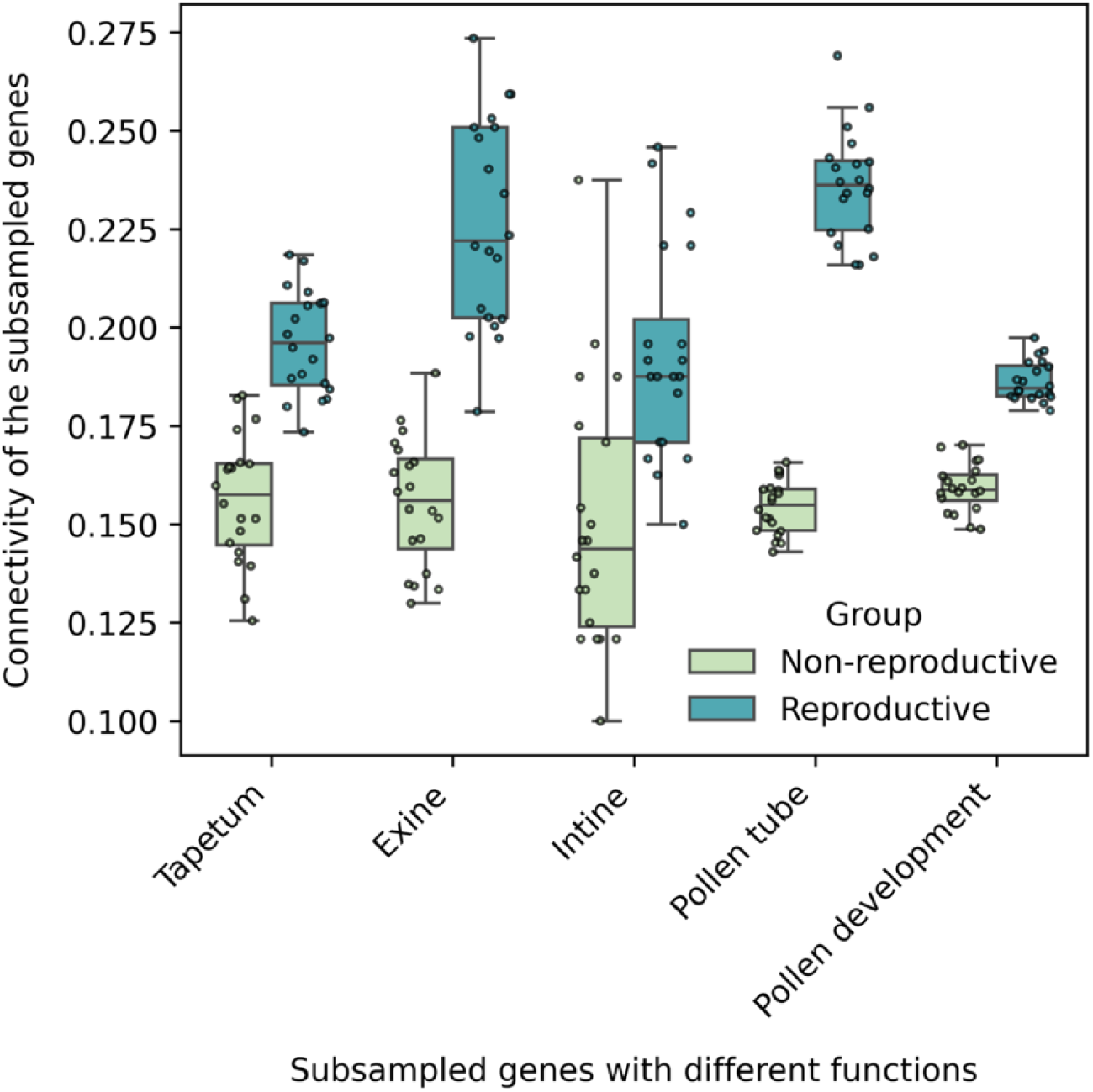
Functional relevance (Connectivity) of pathway genes identified by GO, Mercator, and Plant Connectome. Prior to conserved gene co-expression network (GCN) analysis, genes associated with pollen-related pathways were identified using GO, Mercator, and Plant Connectome annotations. In the *A. thaliana* GCN, functional relevance, i.e., connectivity was evaluated using the score F(S)=mean(si/(|S|−1)), where si represents the number of edges connecting gene *i* to the same pathway genes, and |S| is the total number of genes in the pathway. Edges were defined using a co-expression cutoff of z-score > 1.5. 50% of the pathway genes were randomly subsampled 20 times to generate the figure.

**Figure S7.**
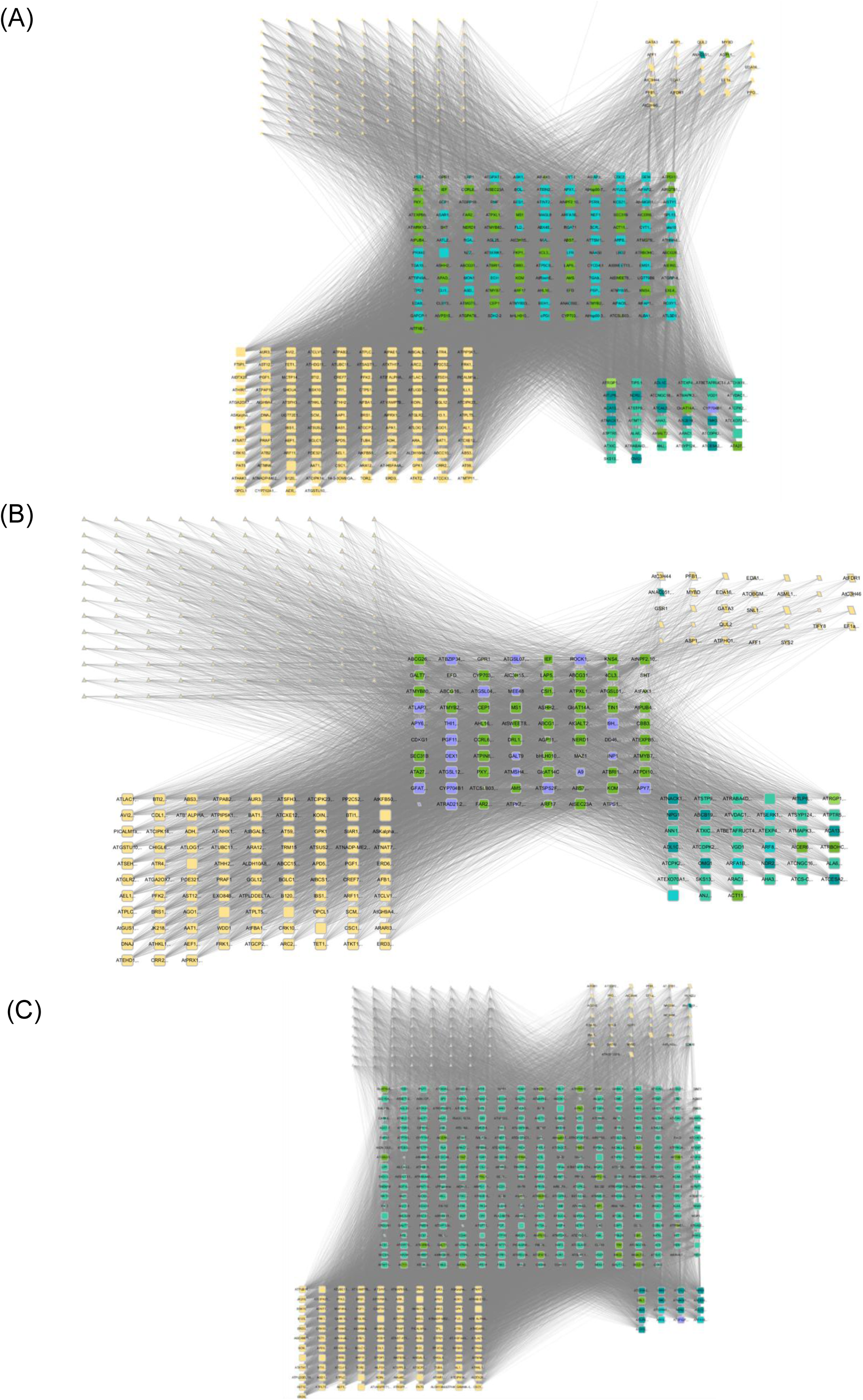
Conserved gene co-expression networks for (A) tapetum, (B) exine, and (C) pollen tube. Node size represents the number of species in which each gene family is conserved.

**Figure S8.**
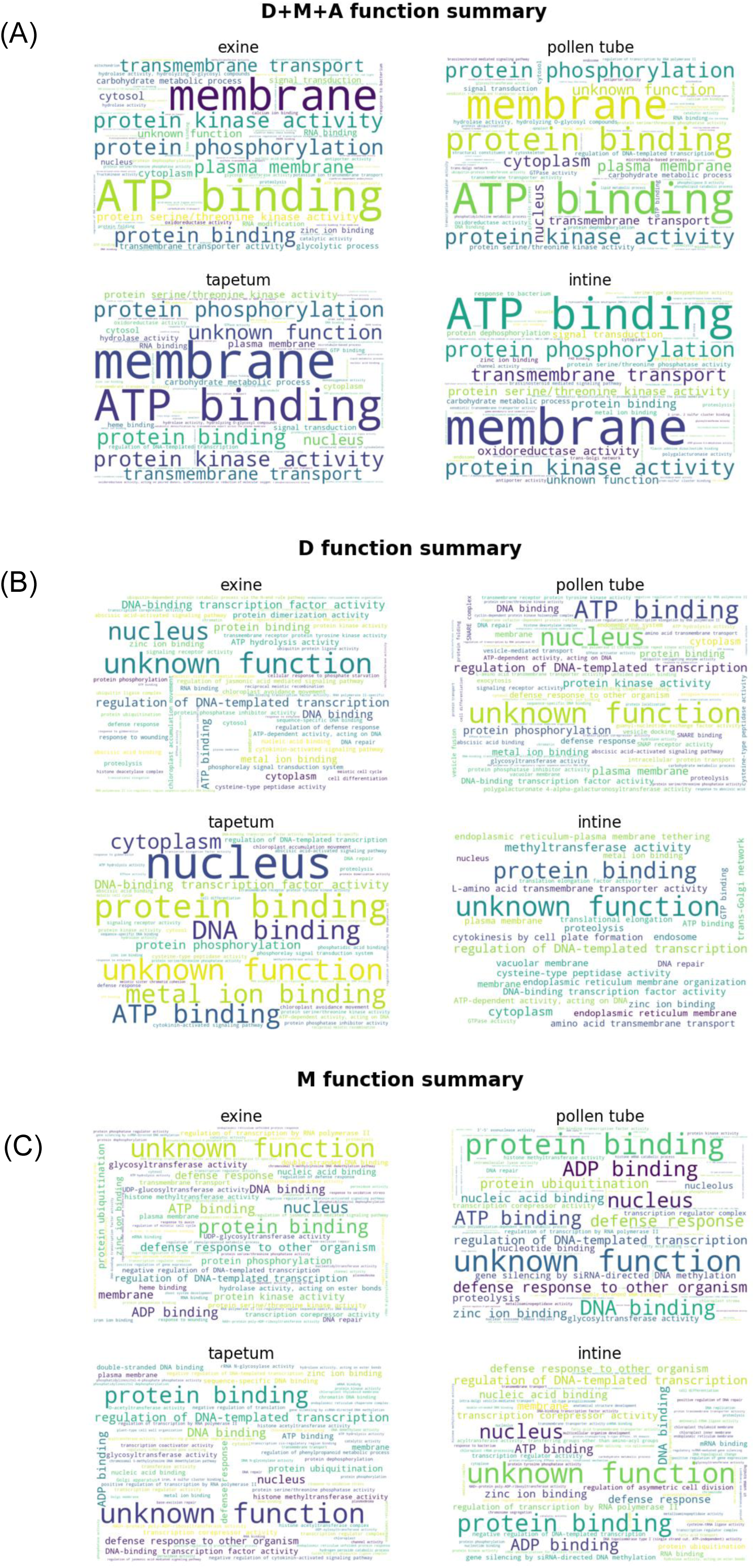
**Word cloud functional summaries of (A) non-reproductive D+M+A (conserved), (B) D (dicot-specifically conserved), and (C) M (monocot-specifically conserved) nodes in exine, pollen tube, tapetum, and intine pathways.**

**Figure S9.**
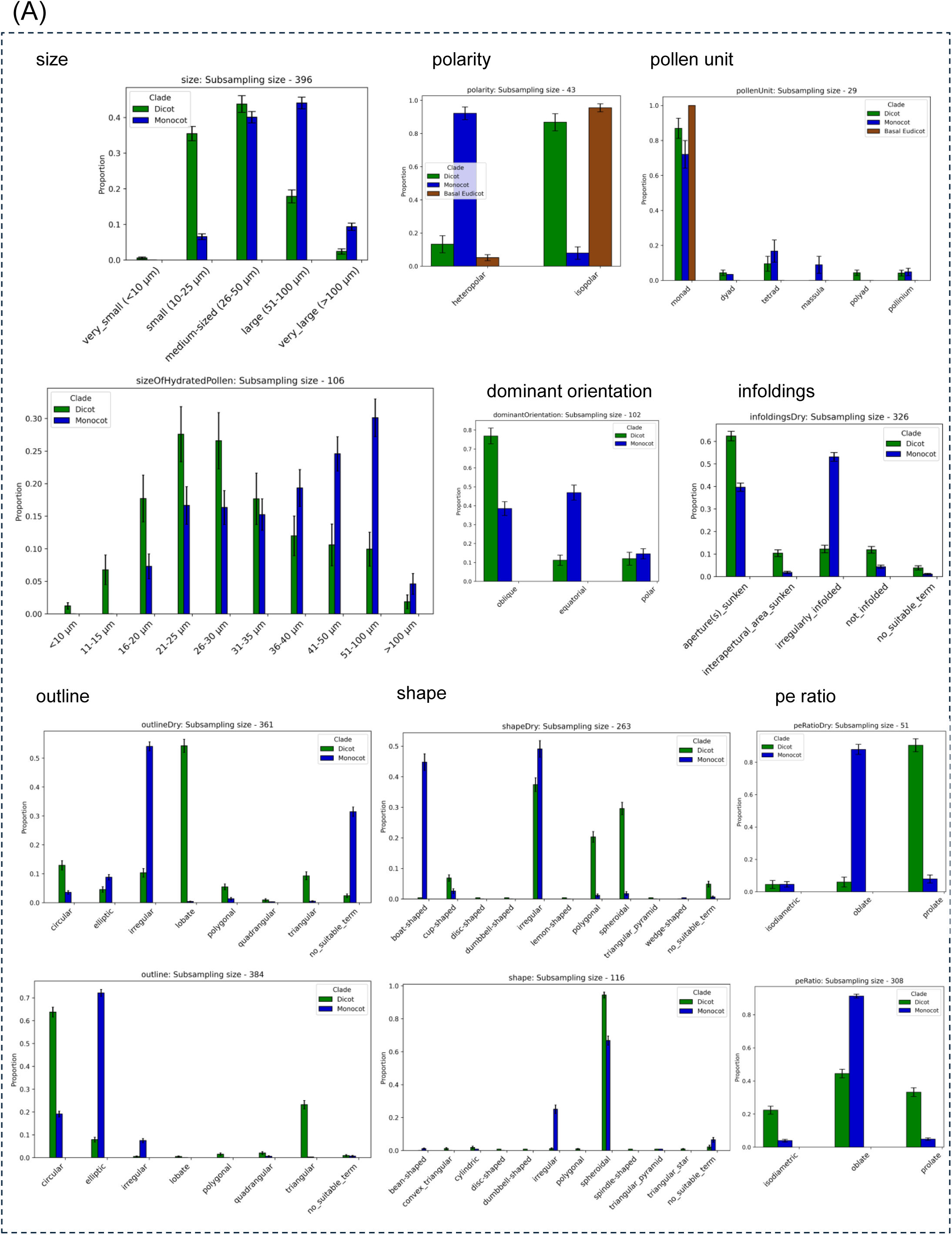
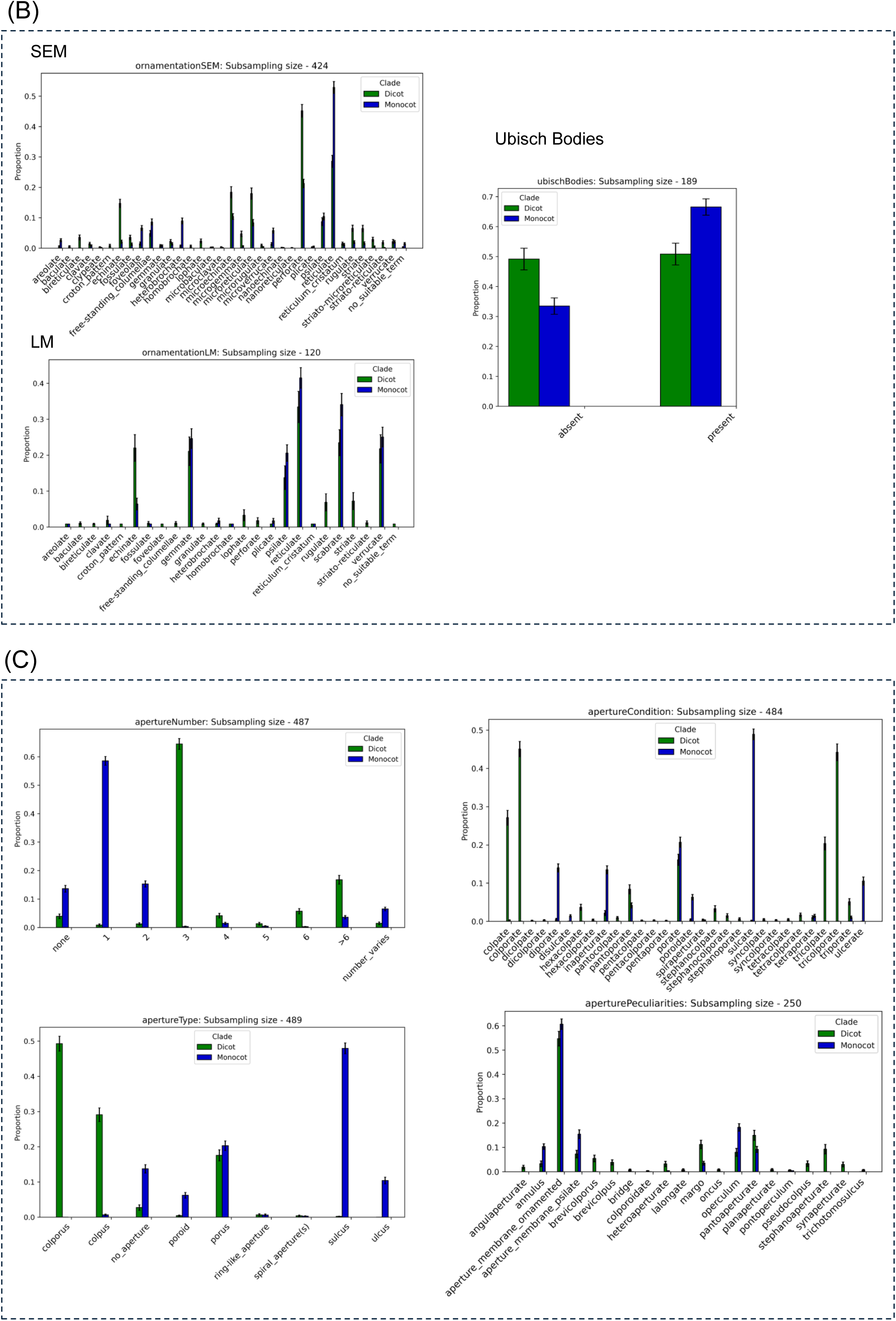
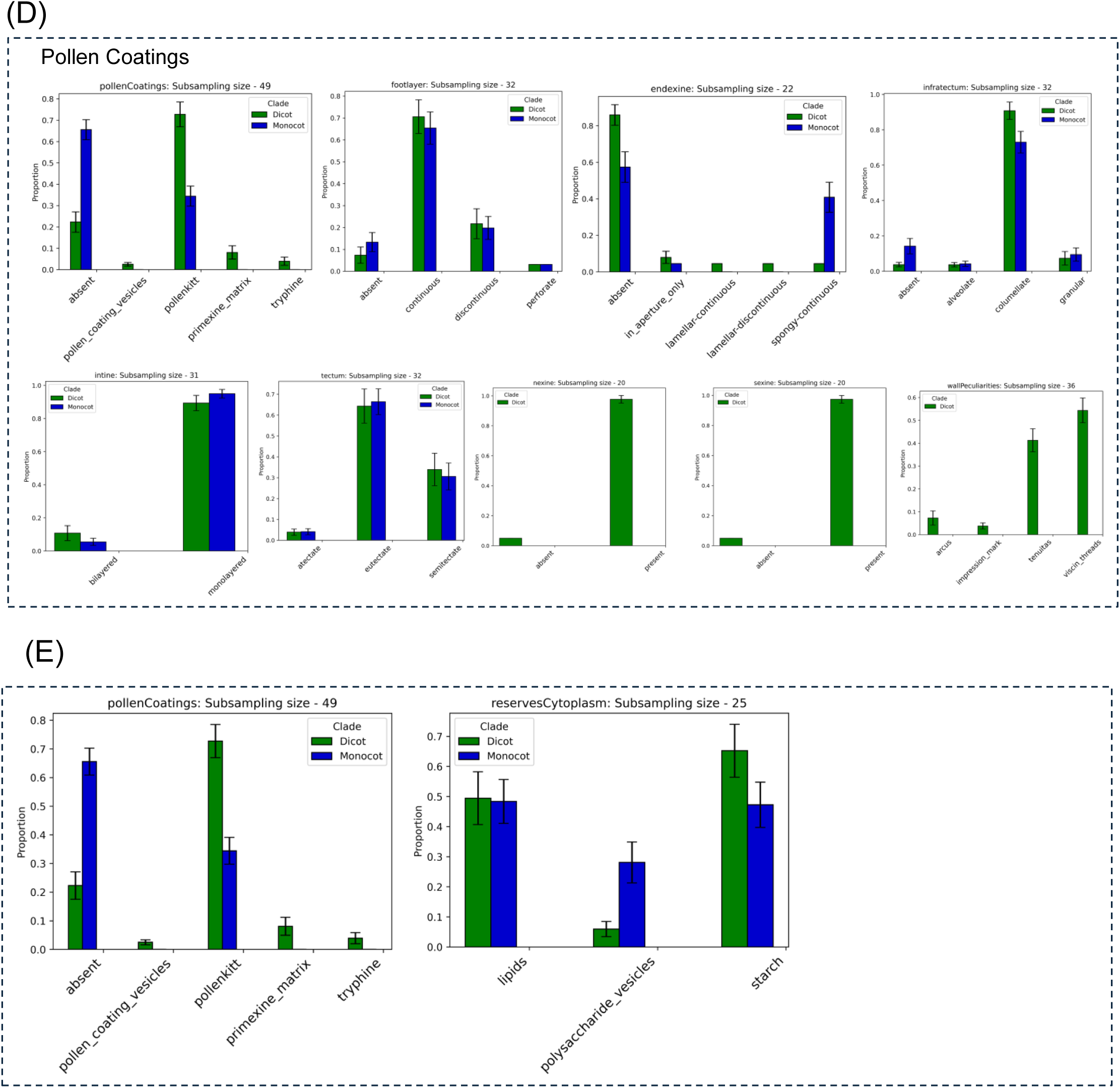
Comprehensive bar graphs from the PalDat database showing differences between dicots (green) and monocots (blue) in pollen (A) overall morphology, (B) surface, (C) **aperture, (D) pollen wall, and (E) cytoplasmic region.**

**Figure S10.**
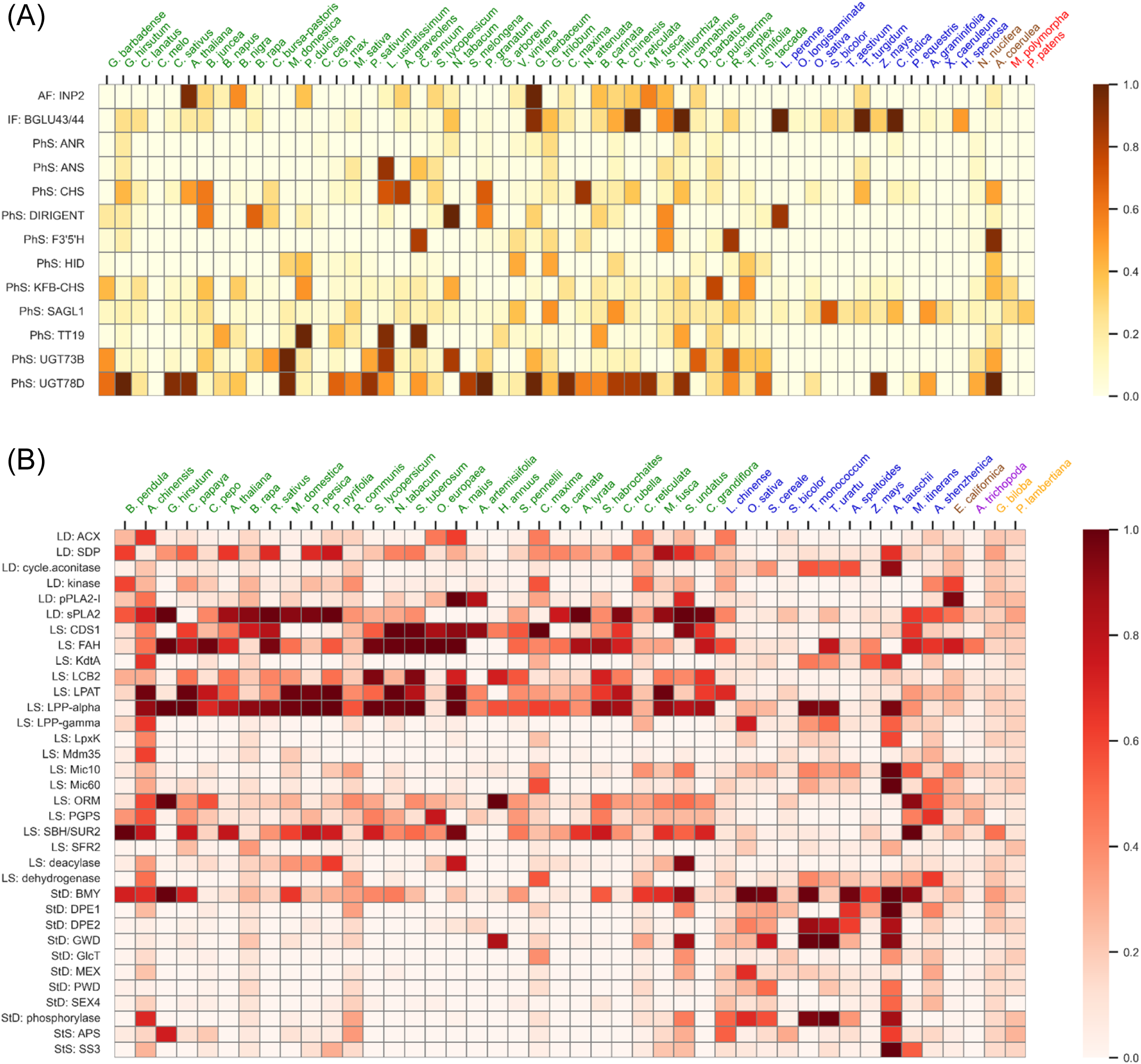
Distribution of gene SPM values across species with anther (top, yellow) and pollen (bottom, red) RNA-seq data.

**Figure S11.**
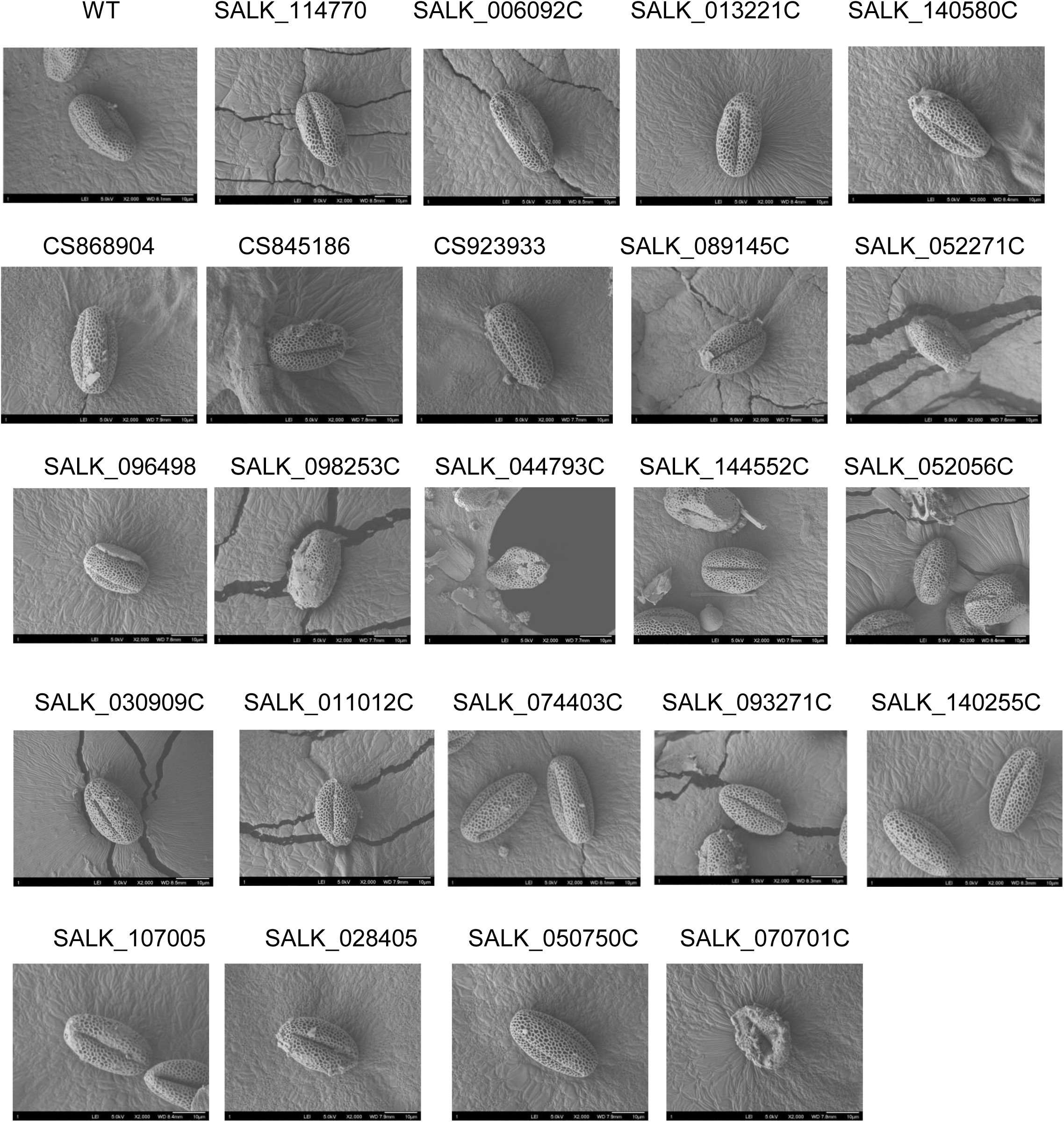
Scanning electron microscopy (SEM) images showing pollen morphology in 20 mutant lines.

**Figure S12.**
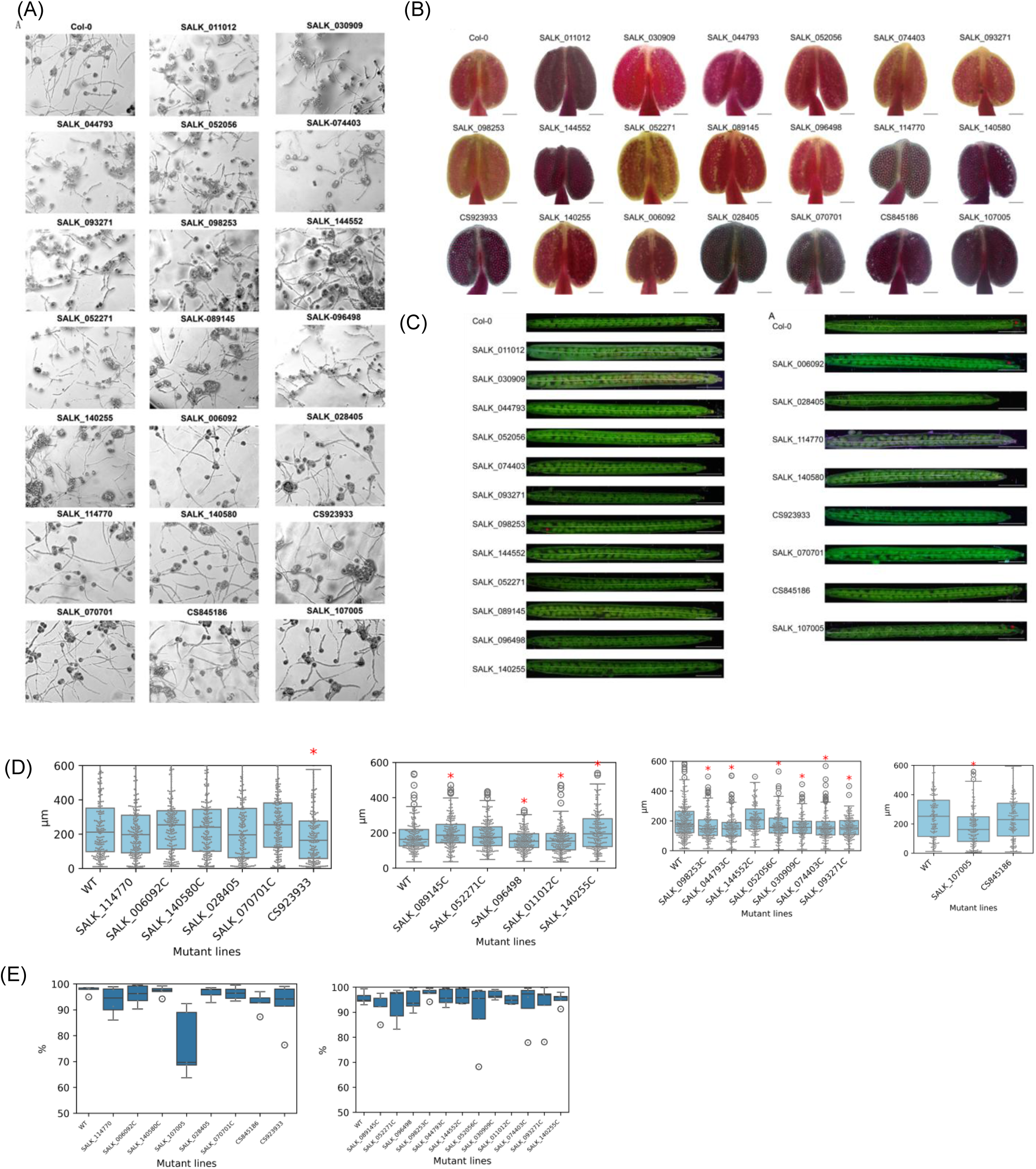
Light microscopy images of *A. thaliana* showing (A) pollen tube, (B) pollen viability within the anther, and (C) siliques from 20 mutant lines. (D,E) Statistical summaries of pollen tube length and pollen germination rate, respectively.

